# Dysregulation of cholesterol and bile acid homeostasis in the gut-liver axis following *Cryptosporidium* intestinal infection

**DOI:** 10.1101/2024.08.01.606233

**Authors:** Shuhong Wang, Ai-Yu Gong, Edward Barker, David L. Williams, Christopher Forsyth, Liqing Yu, Xian-Ming Chen

## Abstract

*Cryptosporidium* spp., an apicomplexan protozoan, is one of the most common pathogens causing moderate-to-severe diarrhea in children under 2 years of age and is also an important opportunistic pathogen for patients with AIDS. There are currently no effective vaccines or therapies available. Infection in children is associated with malnutrition, growth defects, and even impaired cognitive development, but underlying mechanisms remain unclear. We report here that *C. parvum* infection in neonatal mice impairs bile acid reabsorption in the ileum, disrupts lipid metabolism in the liver, and alters bile acid homeostasis in the enterohepatic circulation. The reduction of bile acid pool further impairs lipid absorption in the small intestine. Additionally, replenishing bile prevents the decline in lipid absorption in infected neonatal mice. Strikingly, bile gavage significantly reduces the infection burden and ameliorates the dysregulated homeostasis of cell proliferation and migration in intestinal epithelium following infection. These findings may guide novel therapeutic approaches for cryptosporidiosis.

## Introduction

*Cryptosporidium* is a significant opportunistic intestinal pathogen for immunocompromised individuals and one of the leading causes of moderate-to-severe diarrhea in children under 2 years of age^1^. Infection with this parasite in young kids is associated with mortality, growth stunting, and developmental deficits^2^. Recent estimates from the Global Burden of Diseases, Injuries, and Risk Factors Study (GBD) indicate *Cryptosporidium* spp. is responsible for approximately 7.6 million cases and between 48,000 and 202,000 deaths annually among young children in low-resource settings^3,4^. Cryptosporidiosis in humans is primarily caused by *C. parvum* and *C. hominis* species^2^. Nitazoxanide, the only approved drug for the treatment of cryptosporidiosis, is ineffective in immunocompromised patients and malnourished children^2,5–7^. Pathologically, *Cryptosporidium* infections are primarily confined to the distal ileum and it is speculated that the combined loss of microvillus borders and villus height reduces the absorptive surface area of the intestine, leading to diarrhea and malnutrition in children^8^. However, the current understanding of the underlying molecular mechanisms of malnutrition remains unclear, hampering the development of effective therapeutics. The intestinal mucosa consists of a monolayer of rapidly self-renewing epithelial cells. Intestinal stem cells in the crypts generate new functional epithelial cells, which mature and differentiate as they move up the villus before being shed at the tip^9,10^. *Cryptosporidium* infection is confined to the villi (mainly enterocytes) of the intestinal epithelium, where the organism forms an intracellular yet extra-cytoplasmic parasitophorous vacuole at the brush border following cell entry^11^. Increased cell death during infection triggers crypt hyperplasia and branching due to accelerated cell division to compensate for the loss of cells at the villi^8,12^, however, its underlying molecular mechanisms remain unknown.

As a major source of nutrition for infants, milk provides a significant amount of fatty acids, phospholipids, and cholesterol. Bile acids are amphipathic molecules synthesized from cholesterol in the liver, stored in the gallbladder, and released into the duodenum after a meal^13^. Bile acids facilitate the emulsification of insoluble lipids into tiny mixed micelles, promoting lipid digestion and absorption in the intestine^14^. More than 95% of the bile acids are reabsorbed in the terminal ileum through the apical sodium-dependent bile salt transporter (ASBT; SLC10A2) and recirculated to the liver via the portal blood in a process known as enterohepatic circulation^13^. Bile acids have been identified as endogenous ligands with high affinity for the Farnesoid X receptor (FXR), a ligand-activated transcription factor in the nuclear receptor (NR)-family ^15–17^. Thus, the Bile acid-FXR network plays a crucial role in maintaining whole-body bile acid homeostasis by regulating the expression of target genes involved in both hepatic bile acid synthesis and enterohepatic circulation^18^. Using a murine neonatal infection model, we report that *Cryptosporidium* infection significantly disrupts bile acid and lipid homeostasis in both the intestine and liver. Functionally, bile intervention greatly reduces the infection burden, enhances intestinal lipid absorption, and ameliorates dysregulated intestinal homeostasis. These findings suggest that bile intervention may have therapeutic potential for intestinal cryptosporidiosis.

## Results

### *C. parvum* infection disrupts cholesterol homeostasis in the small intestine

*Cryptosporidium* lacks the capability to synthesize cholesterol and needs to acquire it both from the gut lumen and the host cell to meet its metabolite^19,20^. Therefore, we investigated whether *C. parvum* infection affects intestinal cholesterol absorption and transport. To explore this, we performed RNA-seq analysis to examine genome-wide changes in gene expression in the ileal epithelium of infected neonatal mice. Consistent with results from previous studies^21^, RNA-Seq analysis of the ileal epithelium from infected neonatal mice [5-days old, 72 h post-infection (p.i.)] reveals significant changes in host gene expression (Supplementary Table 1). Gene set enrichment analysis (GSEA) indicates marked alterations in pathways related to inflammatory responses (Extended Data Fig. 2b), defense response (Extended Data Fig. 2c), and the regulation of sterol homeostasis in the infected ileum epithelium (Extended Data Fig. 2d).

Cellular cholesterol homeostasis is finely regulated through the dynamic balance between biosynthesis, uptake, export, and esterification^22^. The regulation of cellular cholesterol transport involves low-density lipoprotein receptor (LDLR)-mediated absorption and chylomicron secretion, as well as transmembrane influx/efflux through transporters such as Niemann-Pick C1-Like 1 (NPC1L1), the heterodimeric ATP-binding cassette transporters G5 and G8 (ABCG5/G8), and ATP-binding cassette transporter A1 (ABCA1) (Fig. 1a). This process is governed by intricate feedback regulation mechanisms, including the liver X receptor (LXR) signaling^23^. Further GSEA analysis indicates that cholesterol efflux, import, and esterase activity were significantly downregulated in the infected ileal epithelium (Fig. 1b and Extended Data Fig. 2a,d), while most genes involved in cholesterol biosynthesis were upregulated (Fig. 1b). Specifically, reduced expression of several cholesterol efflux genes, including ABCG5, ABCG8 and ABCA1, was confirmed by reverse transcription–quantitative PCR (RT–qPCR) (Fig. 1d). Immunoblot analyses confirmed reduced protein levels of ABCG5 and ABCG8 from the ileal epithelium of infected mice (72 h and 96 h p.i.) compared to control animals (Fig. 1f). Two LXRs, LXRα (NR1H3) and LXRβ (NR1H2), play key roles in the transcriptional regulation of genes involved in cholesterol homeostasis^23^. Interestingly, the protein level of LXRβ was decreased, while LXRα level remained unchanged in the ileum of infected neonates compared to the control groups (72 h and 96 h p.i.) (Fig. 1f), suggesting repression of the LXR pathway following infection. In contrast, most cholesterol biosynthesis genes were upregulated after *C. parvum* infection (Fig. 1b,c), as validated by RT-qPCR (Fig. 1e). The rate-limiting enzyme in cholesterol biosynthesis is 3-hydroxy-3-methylglutaryl-CoA reductase (HMGCR)^24^. Immunoblot analyses of cytosolic protein extracts confirmed the higher levels of HMGCR protein in the ileum of infected neonates compared to the control groups (Fig. 1f). Furthermore, we found a significant reduction of free cholesterol in the ileum of infected neonatal mice compared to control mice (Fig. 1h).

**Fig. 1.**
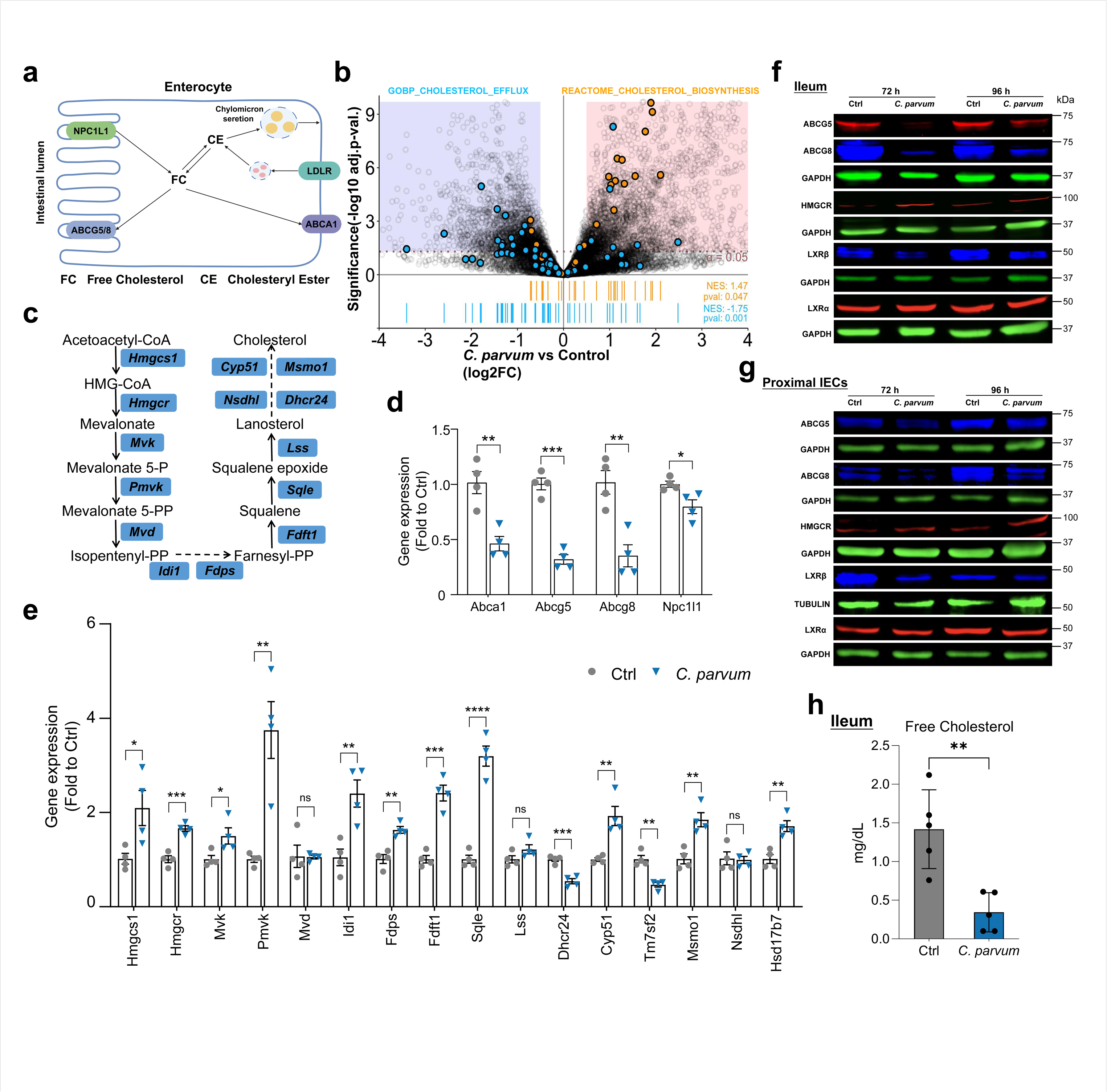
*C. parvum* infection disrupts cholesterol homeostasis in the small intestine. **a,** Schematic diagram summarizing the molecular mechanism of cholesterol transport. **b,** Changes in cholesterol homeostasis-related gene expression upon *C. parvum* infection (72 h p.i.). The enrichment of cholesterol biosynthesis and efflux gene sets is shown at the bottom. The upregulated and downregulated genes (log2FC > 0.5, adj. p < 0.05) are indicated by the light red– and light blue–shaded rectangles underlaid on plot, respectively. **c,** Schematic of the cholesterol biosynthesis pathway, where genes are represented in blue. **d, e,** Relative amounts of mRNAs analyzed by RT–qPCR. Gene expression was presented as fold change to uninfected control. **f, g,** Immunoblotting analysis of ileum and proximal intestinal epithelial cells (IECs) in neonates with and without *C. parvum* infection (72 h and 96 h p.i.). GAPDH and TUBULIN were used as loading control of cytosolic protein. **h,** Quantification of free cholesterol in ileum of control and infected mice (96 h p.i.). Each symbol represents one mouse, all from the same litter. Statistical analysis: unpaired t test for (d, e). Data are expressed as mean ± SEM, and representative of at least two independent experiments, n = 3–5 per group. *P < 0.05, **P < 0.01, ***P < 0.001, ****P < 0.0001. For gel source data, see Supplementary Figure 1.

Strikingly, the proximal small intestine, which is primarily responsible for nutrient absorption and is not infected with *C. parvum* (Extended Data Fig. 1a,b), also exhibited similar changes in the expression levels of cholesterol homeostasis genes (Fig. 1g and Extended Data Fig. 2e). Notably, RT-qPCR showed an increased expression of NPC1L1 in the proximal small intestine but decreased expression in the ileum (Fig. 1d and Extended Data Fig. 2e). Furthermore, a significant reduction in total cholesterol was detected in the proximal small intestinal epithelium of infected neonates compared to the control groups (Fig. 3g and Extended Data Fig. 6a). Most of the cholesterol absorbed by the intestine is secreted as chylomicrons into the circulation^25^. Expression of genes involved in the chylomicron assembly was significantly reduced in both the proximal small intestine and ileum of infected animals (Extended Data Fig. 2f-h). Consistent with results from previous studies^8,21^, the villi length in the infected ileum was significantly shorter (Extended Data Fig. 2j), whereas no such change was observed in the non-directly infected region of the intestine (Extended Data Fig. 2j). Additionally, the small intestine length in infected neonates was shorter compared to control animals (Extended Data Fig. 2i), suggesting a reduction in the absorptive surface area in the intestine. Collectively, these findings indicate that *C. parvum* infection disrupts cholesterol homeostasis in the small intestine.

### *C. parvum* infection alters hepatic lipid metabolism

The liver is the primary organ for cholesterol metabolism, and most lipids in the intestine are absorbed by enterocytes in the small intestine, transported through the lymphatic system into the circulation, then to the liver, and eventually to peripheral tissues^26,27^. Based on this, we hypothesized that intestinal infection by *C. parvum* could impact cholesterol and bile acid metabolism in the liver. To test this, we infected neonatal mice with *C. parvum* oocysts for 3 days, collected liver tissues, and performed RNA-seq (Fig. 2a). Consistent with our hypothesis, *C. parvum* infection caused significant alterations in gene expression profiles in the liver (Extended Data Fig. 3a and Supplementary Table 2). GSEA indicated that the major pathways associated with the altered gene expression profiles are related to immune response regulation (Extended Data Fig. 3a). Gene set variation analysis (GSVA) revealed an overall downregulation of genes involved in lipid metabolism pathways in the liver of infected mice, including these related to cholesterol and bile acid metabolism (Supplementary Table 2). RT-qPCR further validated the reduced levels of most genes involved in lipid metabolism (Extended Data Fig. 3b-f). Immunoblot analyses of APOB100 (Apolipoprotein B100), HMGCR, and HMGCS2 (hydroxymethylglutaryl CoA synthase 2) confirmed reduced protein levels in the infected animals (Extended Data Fig. 3g). The protein levels of hepatic FXR and LXRβ were significantly decreased, whereas LXRα remained unchanged after infection (Fig. 2d). Consistent with this, the mRNA expression of most FXR and LXR target genes was significantly reduced in the liver of infected mice (Fig. 2b,c). Immunoblot of selected canonical target genes of LXRs and FXRs, including *ABCG5*, *ABCG8*, *CYP7A1* and *NR0B2* (*Shp*), confirmed decreased protein levels (Fig. 2d). Accordingly, lipidomics analysis revealed altered lipid profiles in the liver of infected mice (Fig. 2a and Extended Data Fig. 4a,b). There was no significant reduction in total lipids in the liver after 4 days infection compared to the control group (Extended Data Fig. 4d,e). However, reduced amounts of bile acids and cholesterol esters, but not total cholesterol, were detected in the liver of infected mice (Fig. 2e and Extended Data Fig. 4f-h). As an important component of milk, fat provides about 50% of the energy for infants^28^. The main fat component is triacylglycerol (TG), which makes up more than 98% of milk fat^29^. We observed significant reductions in most TGs, diacylglycerols (DG), and free fatty acids (FFA) in the liver after *C. parvum* intestinal infection (Fig. 2e and Extended Data Fig. 4i-n). Kyoto Encyclopedia of Genes and Genomes (KEGG) pathway enrichment analysis of the lipidomics results revealed downregulation of lipid metabolic pathways in the liver of infected mice (Extended Data Fig. 4c), consistent with the GSVA results from the RNA-seq (Supplementary Table 2). To further investigate the alterations in bile acids, we performed a comprehensive, quantitative bile acid targeted assay (Fig. 2a), which revealed broad metabolic downregulation in the liver of infected mice (Fig. 2f-h). Taken together, these data suggest that *C. parvum* intestinal infection disturbs cholesterol and bile acid homeostasis in the liver.

**Fig. 2.**
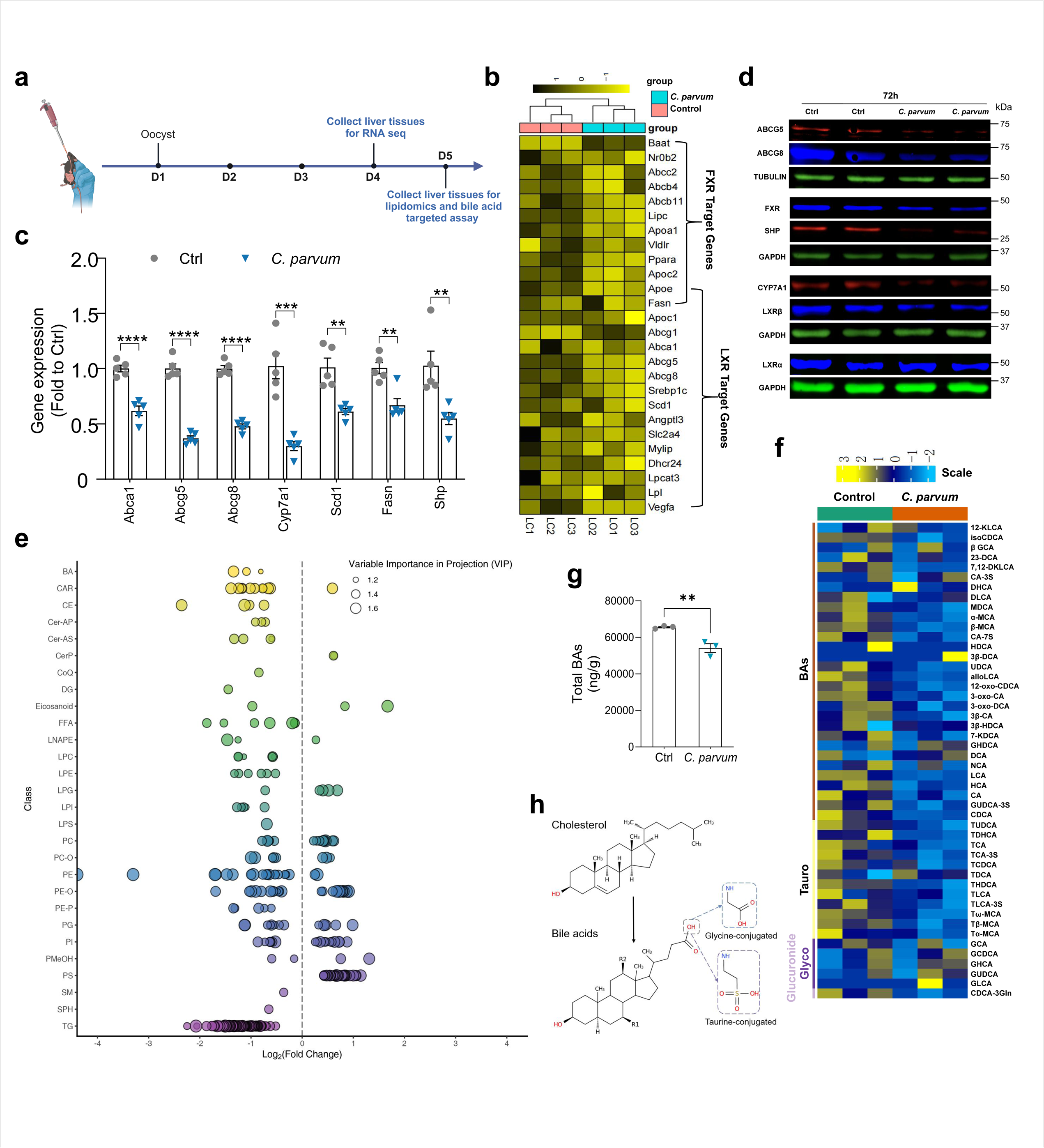
*C. parvum* infection alters hepatic lipid metabolism. **a,** Experiment schematics: Neonatal mice (5 days old) were orally inoculated with *C. parvum* oocysts (10^6^ oocysts per animal), and liver tissues were collected for deep sequencing (72 h p.i.) and metabolomics (96 h p.i.). **b,** Clustered heatmap of normalized gene counts of LXR and FXR target genes expression in the liver of *C. parvum* infected neonatal mice versus control mice (72 h p.i.). **c,** RT-qPCR analysis confirming hepatic mRNA expression of LXR and FXR target genes in neonates with and without *C. parvum* infection at 72 h p.i. **d,** Immunoblotting analysis of liver samples in control and *C. parvum* infected neonatal mice (72 h p.i.). GAPDH and TUBULIN were used as loading control of cytosolic protein. **e,** Scatter plot showing the change of hepatic lipids between control or *C. parvum* infected mice in different subclasses (96 h p.i.). Lipid with VIP ≥ 1 represents significant difference. **f,** Heatmap of differential hepatic bile acids expression in control versus *C. parvum* infected mice (96 h p.i.). **g,** The total quantification of all bile acids in liver between control and *C. parvum* infected neonatal mice from (f). **h,** Structures of cholesterol and bile acids. Bile acids are synthesized from cholesterol and can be amidated by taurine (or glycine). Each symbol represents one mouse, all from the same litter. Statistical analysis: unpaired t test for (c, g). Data are expressed as mean ± SEM, and representative of at least two independent experiments, n = 3–5 per group. *P < 0.05, **P < 0.01, ***P < 0.001, ****P < 0.0001. For gel source data, see Supplementary Figure 1.

### *C. parvum* infection decreases bile acid levels in the small intestine and reduces the absorption of dietary lipids

Given the significant dysregulation of cholesterol and bile acid homeostasis in the liver following *C. parvum* intestinal infection, particularly the decreased expression of FXR-targeted genes, we next investigated lipid metabolism with a focus on the Bile acid-FXR signaling pathway in the infected intestine. FXR is essential for maintaining whole-body bile acid homeostasis by regulating the expression of target genes involved in both hepatic bile acid synthesis and enterohepatic circulation^18^. The enterohepatic circulation of bile acids and the FXR pathway involve the synthesis of bile acids from cholesterol in the liver and the finely controlled expression of genes for bile acids reabsorption in the intestinal epithelium, as shown in Fig. 3a. GSEA indicates that many genes involved in the bile acid metabolic process in the ileum were significantly reduced in infected mice (Fig. 3b). The reduction of FXR and FXR-targeted genes *Shp* and *Fgf15* suggests a robust suppression of FXR signaling in the small intestine following infection (Fig. 3b-d and Extended Data Fig. 5a). Bile acids generate negative feedback on ileal ASBT transcription through FXR-mediated induction of *Shp*^30^. Notably, despite a substantial increase in its mRNA expression (Fig. 3b,c), ASBT was markedly reduced at the protein level (Fig. 3d), which could substantially impact bile reabsorption in the ileum.

**Fig. 3.**
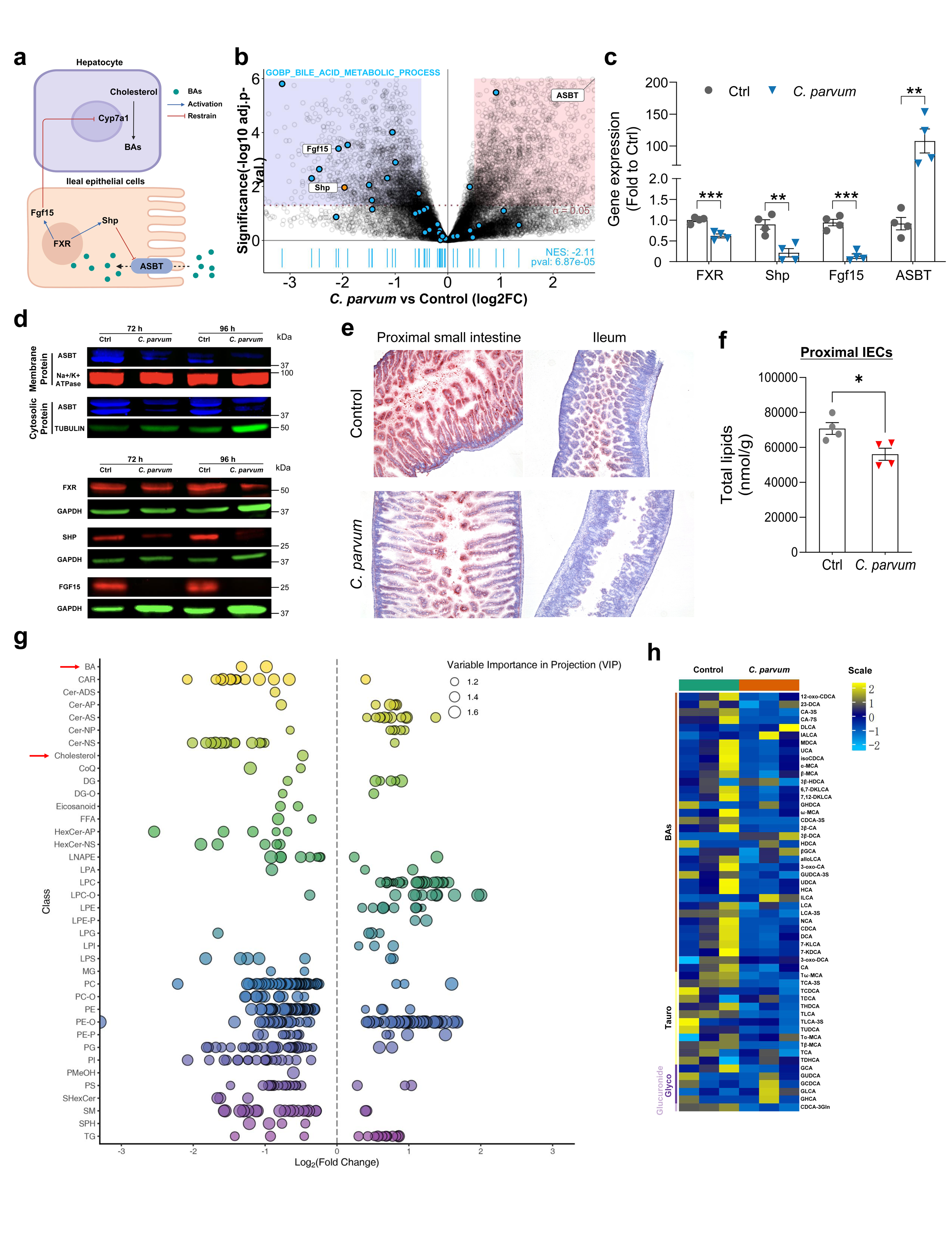
*C. parvum* infection decreases bile acid levels in the small intestine and reduces the absorption of dietary lipids. **a,** Schematic overview of the enterohepatic circulation of bile acids and the FXR pathway. **b,** Differential gene expression analysis upon *C. parvum* infected neonatal mice (72 h p.i.). The enrichment of bile acid metabolic gene set is shown in blue and the distribution of Shp and ASBT is shown in orange. **c,** RT-qPCR analysis confirming ileum mRNA expression of FXR, Fgf15, Shp and ASBT from infected neonates (72 h p.i.). **d,** Immunoblotting analysis of ileum samples in control and *C. parvum* infected neonatal mice (72 h and 96 h p.i.). GAPDH and TUBULIN were used as loading control of cytosolic protein, Na+/K+ ATPase was used as loading control of membrane protein. **e,** Oil Red O stains of the proximal small intestine and ileum section in neonates with and without *C. parvum* infection (96 h p.i.). **f,** The total quantification of all lipids in proximal IECs between control and *C. parvum* infected neonatal mice. **g,** Scatter plot showing the change of proximal IECs lipids between control or *C. parvum* infected mice in different subclasses. Neonatal mice (5 days old) were orally inoculated with *C. parvum* oocysts (10^6^ oocysts per animal) and proximal IECs were collected (96 h p.i.) for lipidomics analysis. Lipid with VIP ≥ 1 represents significant difference. **h,** Heatmap of differential intestinal bile acids expression in control versus *C. parvum* infected mice. Neonatal mice (5 days old) were orally inoculated with *C. parvum* oocysts (10^6^ oocysts per animal) and small intestine was collected (96 h p.i.) for bile acid targeted assay. Each symbol represents one mouse, all from the same litter. Statistical analysis: unpaired t test for (c, f). Data are expressed as mean ± SEM, and representative of at least two independent experiments, n = 3–4 per group. *P < 0.05, **P < 0.01, ***P < 0.001, ****P < 0.0001. For gel source data, see Supplementary Figure 1.

To address whether dysregulation of liver and intestinal bile acid metabolism affects the bile levels in the small intestine following infection, we infected neonatal mice with *C. parvum* oocysts for 4 days. We then collected small intestine tissues, including the luminal contents, and performed a comprehensive, quantitative bile acid targeted assay. A marked decrease of most intestinal bile acids was observed in the infected mice (Fig. 3h and Extended Data Fig. 6d). In addition, the ileum appears paler, possibly due to a reduction in intestinal bile acids following the infection (Extended Data Fig. 5d). RNA-seq analysis of ileal tissues revealed a global decrease in the expression of genes involved in lipid and vitamin absorption in the infected mice (Extended Data Fig. 5g,h). Imaging of neutral lipids by oil red O (ORO) staining^31^ showed a reduction in neutral lipids in the proximal small intestine and ileum following infection (Fig. 3e). Consistent with this, lipidomics analysis demonstrated a significant decrease in total lipids (Fig. 3f), particularly bile acid, cholesterol, phosphatidylcholines (PC), hexosylceramides (HexCer-NS) and Ceramide-NS (Cer-NS) in the proximal small intestinal epithelium of infected neonates (Fig. 3g and Extended Data Fig. 6a). However, lipidomics analysis showed no significant alteration in total TG level (Fig. 3g and Extended Data Fig. 6a). Additionally, the proximal small intestinal epithelium and ileum from infected neonates exhibited reduced expression of several genes involved in fatty acid oxidation, TG catabolism (Extended Data Fig. 5b,c,e,f), and chylomicron assembly (Extended Data Fig. 2f-h). Furthermore, lower levels of L-palmitoylcarnitine were detected in the proximal small intestinal epithelium of *C. parvum* infected neonates (Extended Data Fig. 6c), indicating impaired fatty acid oxidation. These alterations may contribute to the accumulation of TG in the proximal small intestinal epithelium following infection. Linoleic acid is an essential lipid that cannot be synthesized in mammalian cells and is exclusively acquired from the diet^32^. The KEGG pathway enrichment analysis of lipidomics revealed a reduction in linoleic acid metabolism (Extended Data Fig. 6b), further supporting the reduced absorption of dietary lipids in the intestine after *C. parvum* infection.

### Bile intervention shows therapeutic potential for intestinal cryptosporidiosis

To assess the impact of reduced bile on the pathogenesis of *C. parvum* infection, we administered whole bile collected from the gallbladders of adult mice to neonates (5 days old) at 24 h after oral gavage of *C. parvum* oocysts, followed by daily administration of bile for an additional 3 days (Fig. 4a). Bile feeding activated intestinal transcription of FXR target genes (Fig. 4c) and surprisingly, greatly reduced the infection burden compared to the animals receiving PBS (Fig. 4b). Our data revealed significant variation in infection burden across different regions, with the highest levels typically observed in the distal ileum (Fig. 4d). Notable reductions in the infected area of the distal ileum and overall reduced infection burden were observed in the intestinal tissues of animals receiving bile, compared to the PBS-treated control group (Fig. 4d). Additionally, replenishing bile restored the ileal expression of genes involved in cholesterol homeostasis in the infected mice (Extended Data Fig. 7a,b). Furthermore, human intestinal epithelial HCT-8 cells were first exposed to *C. parvum* infection for 4 h (a time frame for parasite attachment and invasion)^33^, after which free-parasites were removed after washing. Cell cultures were then exposed to various concentrations of bile or PBS for an additional 20 h, followed by measurement of infection burden. A substantial reduction in infection burden was observed in cells treated with higher concentrations of bile (0.5% and 1%) compared to lower bile concentrations (0.2%, 0.25% and 0.33%) (Fig. 4e,f). Interestingly, an increased infection burden was noted in cultures treated with lower concentrations of bile compared to the PBS group (Fig. 4e,f). Primary 2D intestinal epithelial monolayers derived from mice for *C. parvum ex vivo* infection further confirmed this phenomenon (Extended Data Fig. 7e). Data from previous reports show that bile salt sodium taurocholate enhances excystation and the entry of *Cryptosporidium* into host cells^34^. Consistent with this, we observed an overall increase in infection burden when HCT-8 cells were exposed to *C. parvum* oocysts in the presence of various bile concentrations (from 0.07%-2% bile acids as tested) during the 4 h parasite attachment and invasion time frame (Extended Data Fig. 7d). HCT-8 cells exposed to different concentrations of bile, with or without *C. parvum* infection, showed no significant reduction in cell viability (Extended Data Fig. 8a-d). To further assess the therapeutic potential of bile for cryptosporidiosis. We orally inoculated neonatal mice with excessive dose of *C. parvum* (2×10^6^ oocysts per animal). At 24 h after *C. parvum* infection, mice were fed different doses of whole bile collected from the gallbladders of adult mice (0.5, 1.5, and 3 μl of bile per animal, diluted with PBS), followed by daily for another 3 days (Fig. 5a). Resultant data further support a role for bile in reducing infection and increasing lipid uptake following *C. parvum* infection (Fig. 5b-f and Extended Data Fig. 7c). Specifically, feeding 1.5 and 3 μl of bile per day for 4 days activated intestinal transcription of FXR target genes (Fig. 5e) and greatly decreased the infection burden compared to the animals receiving PBS (Fig. 5d,f and Extended Data Fig. 7c), while feeding 0.5 μl of bile per day did not cause significant changes (Fig. 5b,c ). Notably, in the presence of excessive dose of *C. parvum* inoculation, treatment with 3 μl of bile per day for 4 days greatly reduced the infected area from distal ileum in mice, which was negatively correlated with the area and extent of intestinal lipid absorption (Fig. 5f and Extended Data Fig. 7c).

**Fig. 4.**
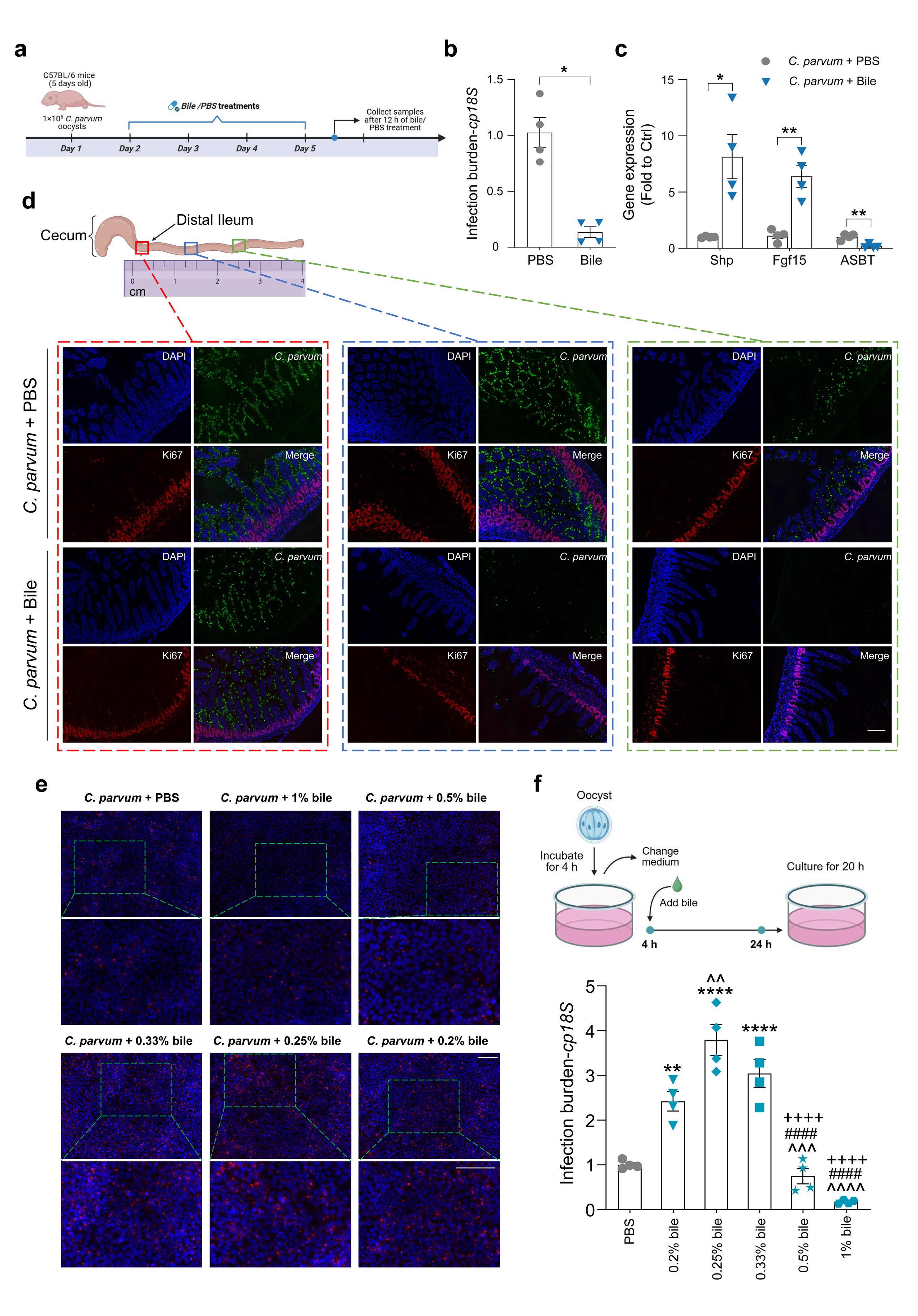
Bile intervention promotes intestinal anti-*C. parvum* defense. **a,** Experiment schematics: Neonatal mice (5 days old) were orally administered 1×10^5^ *C. parvum* oocysts; PBS and bile were given at 24 h p.i. followed by daily for 4 days, respectively; Intestinal tissues were collected, and the mRNA levels of infection burden (b) and FXR target genes (c) were measured by RT-qPCR. **d,** Immunofluorescent staining of *C. parvum* from (a). A higher magnification of the boxed region to visualize the infection burden of *C. parvum* from different portions of ileum is shown (d). Blue: DAPI, red: Ki67, green: *C. parvum*. Bars, 100 µm. **e,** Effects of bile on *C. parvum* infection of cultured HCT-8 cells. Cells were incubated with oocysts for 4 h first, subsequently exposed to different concentrations of bile or PBS for another 20 h, followed by immunofluorescence staining of *C. parvum*. Blue: DAPI (nuclei), red: *C. parvum*. Bars, 100 µm. **f,** Effects of bile on *C. parvum* infection of cultured HCT-8 cells. Cells were incubated with oocysts for 4 h first, subsequently exposed to different concentration of bile or PBS for another 20 h, followed by RT-qPCR of *cp18S* (*p<0.05 vs. PBS, ^ p<0.05 vs. 0.2% bile, #p<0.05 vs. 0.25% bile, + p<0.05 vs. 0.33% bile). Each symbol represents one mouse, all from the same litter. Statistical analysis: unpaired t test for (b, c), one-way analysis of variance (ANOVA) with Tukey tests was used for correction of multiple comparisons in f. Data are expressed as mean ± SEM, and representative of at least two independent experiments, n = 4 per group. *P < 0.05, **P < 0.01, ***P < 0.001, ****P < 0.0001.

**Fig. 5.**
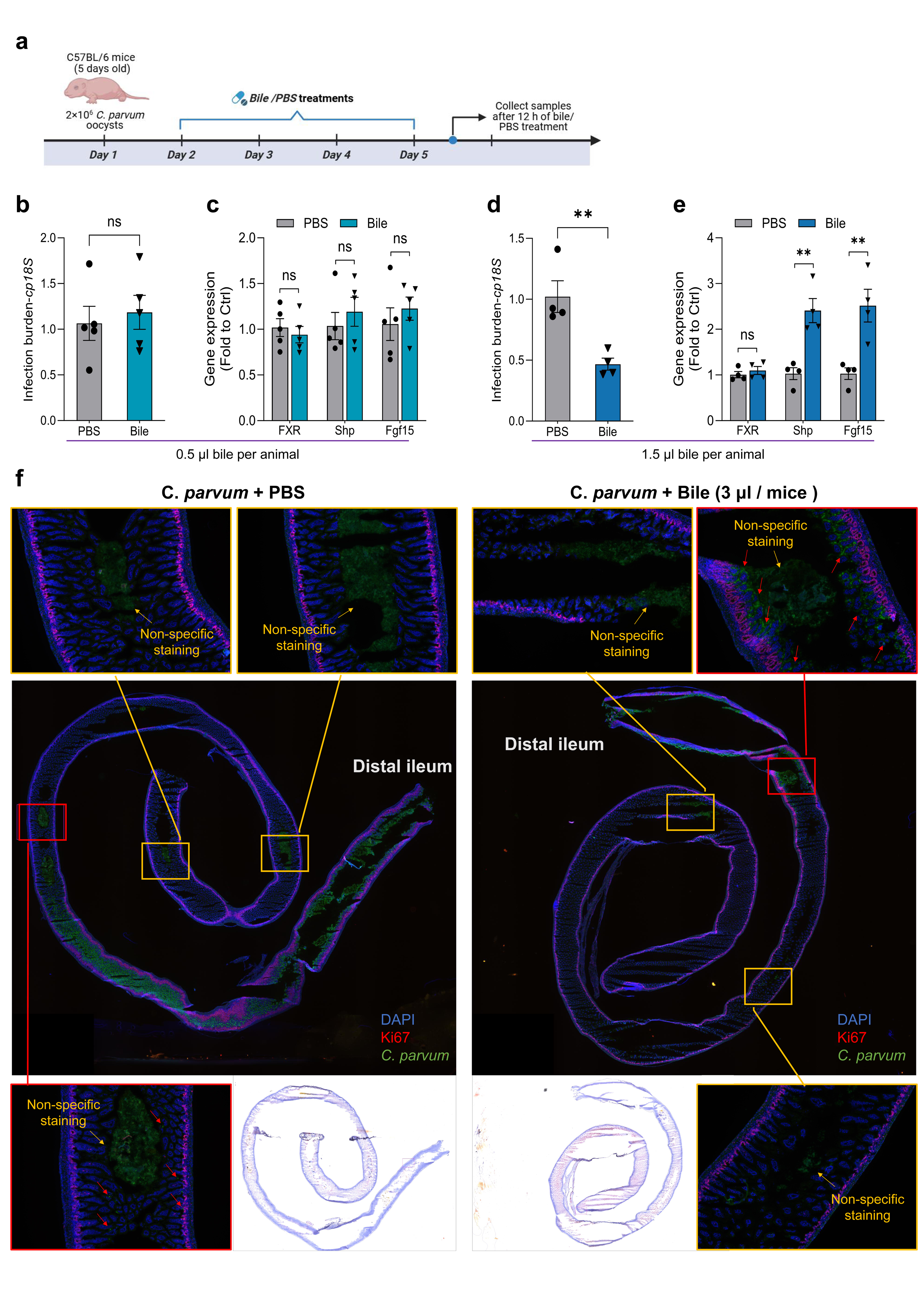
Effects of different doses of bile treatment on *C. parvum* infection *in vivo*. **a,** Experiment schematics: Neonatal mice (5 days old) were orally administered 2×10^6^ *C. parvum* oocysts; PBS and bile were given at 24 h p.i. followed by daily for 4 days, respectively; Intestinal tissues were collected, and the mRNA levels of infection burden (b, d) and FXR target genes (c, e) were measured by RT-qPCR. **f,** Immunofluorescent and Oil Red O staining of the distal small intestine sections from (a). A higher magnification of the boxed region (orange) to visualize the background staining; A higher magnification of the boxed region (red) to visualize the *C. parvum* staining; Red arrow indicates *C. parvum* staining. Blue: DAPI, red: Ki67, green: *C. parvum*. Statistical analysis: unpaired t test for (b-e). Data are expressed as mean ± SEM, and representative of at least two independent experiments, n = 4-5 per group. *P < 0.05, **P < 0.01, ***P < 0.001, ****P < 0.0001.

### Bile acid-FXR signaling pathway is involved in the regulation of intestinal epithelium turnover in response to *C. parvum* infection

To monitor the cell proliferation and epithelial turnover in the intestine following infection, we examined the incorporation of ethynyl-2’-deoxyuridine (EdU) and tracked the migration of EdU-labeled cells at 4 and 48 h after EdU pulse labeling (Fig. 6a). To explore the potential effects of bile, we gavaged infected and uninfected mice with the whole bile collected from the gallbladders of adult mice (Fig. 6a). As shown in Fig. 6e and Extended Data Fig. 9a, there was a significant increase in the number of EdU-positive proliferating cells in the crypt regions of infected mice at 4 h. These proliferative cells also migrated much higher along the villus compared to their control counterparts at 48 h p.i. (Fig. 6g,j). Interestingly, quantification showed an appreciable decrease in EdU-positive proliferating and migrating cells in infected mice treated with bile compared to those treated with PBS (Fig. 6e,g,j and Extended Data Fig. 9a). Notably, we found that the dysregulation of cell proliferation and migration induced by infection, as well as the effects of bile supplement, were not limited to the ileum but extended to the entire small intestine and even the colon (Fig. 6c-j and Extended Data Fig. 9a).

**Fig. 6.**
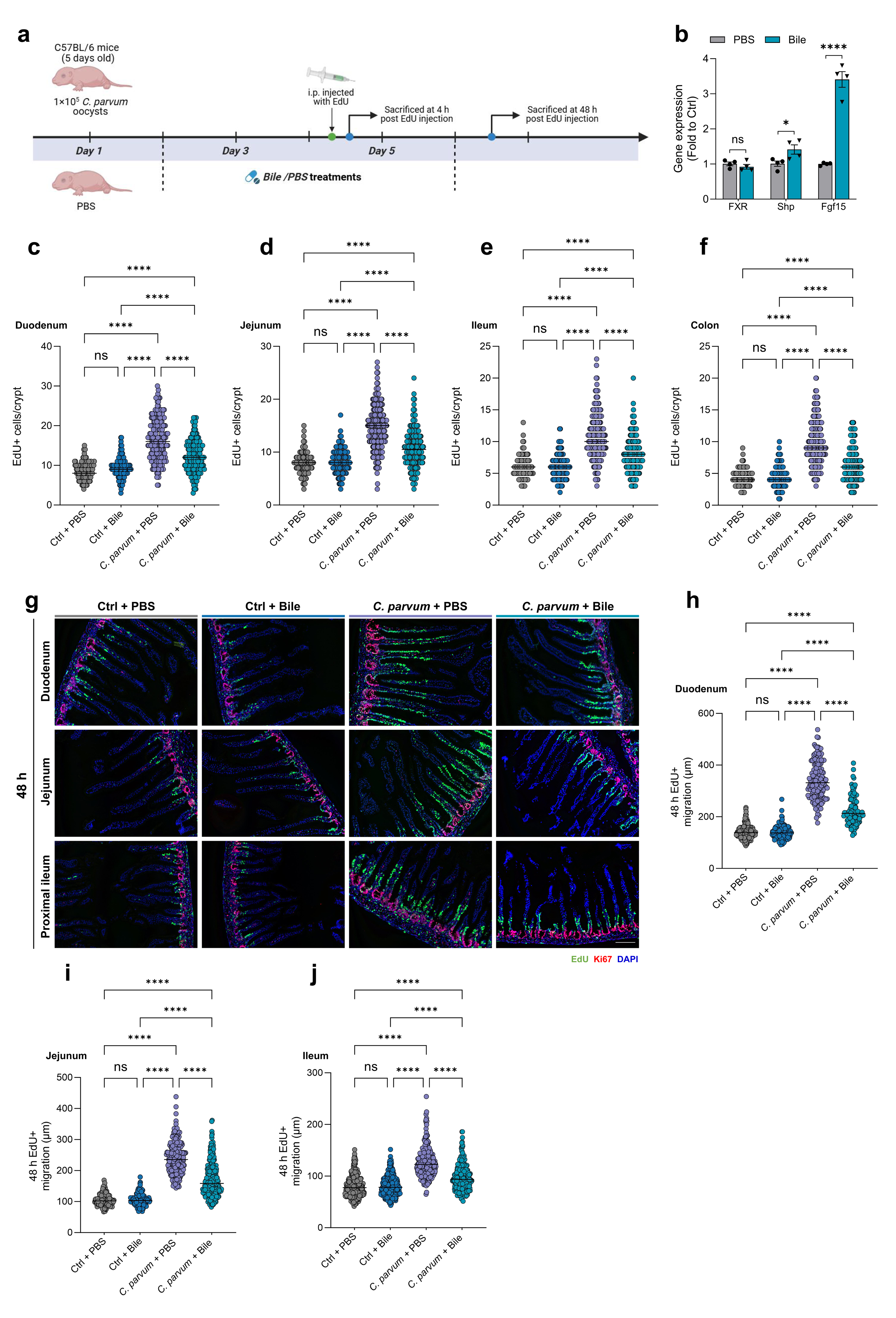
Bile acid treatment attenuates the dysregulated homeostasis in the intestinal epithelium following *C. parvum* infection. **a,** Experiment schematics: Neonatal mice (5 days old) were orally administered 1×10^5^ *C. parvum* oocysts; PBS or bile was given at 24 h p.i. followed by daily for 3 days; Mice were i.p. injected with EdU and sacrificed at 4 h post injection; Mice were i.p. injected with EdU, treated with PBS or bile for another 2 days and sacrificed at 48 h post injection. **b,** RT-qPCR analysis confirming mRNA expression of FXR, Shp, and Fgf15 in the small intestine of uninfected neonates after bile treatment. **c-f**, Quantification of EdU staining of proliferating cells in crypts at 4 h post injection (>50 crypts per mouse, 3 mice/group). **g,** Representative images of Ki67, EdU staining of proliferating cells at 48 h post injection. Blue: DAPI (nuclei), red: Ki67, green: EdU. Bars, 100 µm. **h-j,** Quantification of cell migration length along the crypt-villus axis at 48 h post injection. Quantification of cell migration length by measuring the distance from the crypt base to the highest labelled cell in the villus (>30 villus per mouse, 3 mice/group). Statistical analysis: unpaired t test for (b); One-way analysis of variance (ANOVA) with Tukey tests was used for correction of multiple comparisons in c-f, h-j, and representative of at least two independent experiments, n = 3 per group. *P < 0.05, **P < 0.01, ***P < 0.001, ****P < 0.0001.

There was a significant increase in the transcription levels of FXR target genes in infected mice treated with bile compared to those treated with PBS (Fig. 4c and Fig. 5e). We then examined whether activation of FXR pathway could attenuate crypt hyperproliferation and cell migration *in vivo*. We administrated mice with Fexaramine (FexD) (Fig. 7a), which has previously been shown to activate FXR pathways *in vivo* ^35^. Quantification showed a significant decrease in EdU-positive proliferating and migrating cells (jejunum and ileum) in uninfected mice treated with FexD compared to those treated with vehicle control (Fig. 7b-g). This was consistent with previous literature that the activation of FXR pathway inversely correlates with epithelial proliferation, colorectal cancer progression and malignancy ^36–38^. Interestingly, EdU-labeling analysis demonstrated an even more pronounced decrease in EdU-positive proliferating and migrating cells between infected mice treated with FexD and vehicles, compared to those uninfected mice treated with FexD and vehicles (Fig. 7b-g). Notably, FXR activation significantly increased the infection burden (Fig. 7h,i), further emphasizing the critical role of FXR signaling in maintaining crypt-villus homeostasis during *Cryptosporidium* infection. Importantly, this effect appears to be independent of the infection burden. RT-PCR indicated that bile treatment for 6 h can activate intestinal transcription of FXR target genes in uninfected mice (Fig. 6b). However, in contrast to *C. parvum* infected mice, no significant differences were observed in uninfected mice treated with bile and PBS (Fig. 6c-j and Extended Data Fig. 9a). The negative feedback of bile acid-FXR axis might restrain the constant activation of FXR pathway in uninfected mice ^18^. Overall, these data suggest that the Bile acid-FXR signaling pathway is involved in regulating cell proliferation and migration of the intestinal epithelium following *C. parvum* infection.

**Fig. 7.**
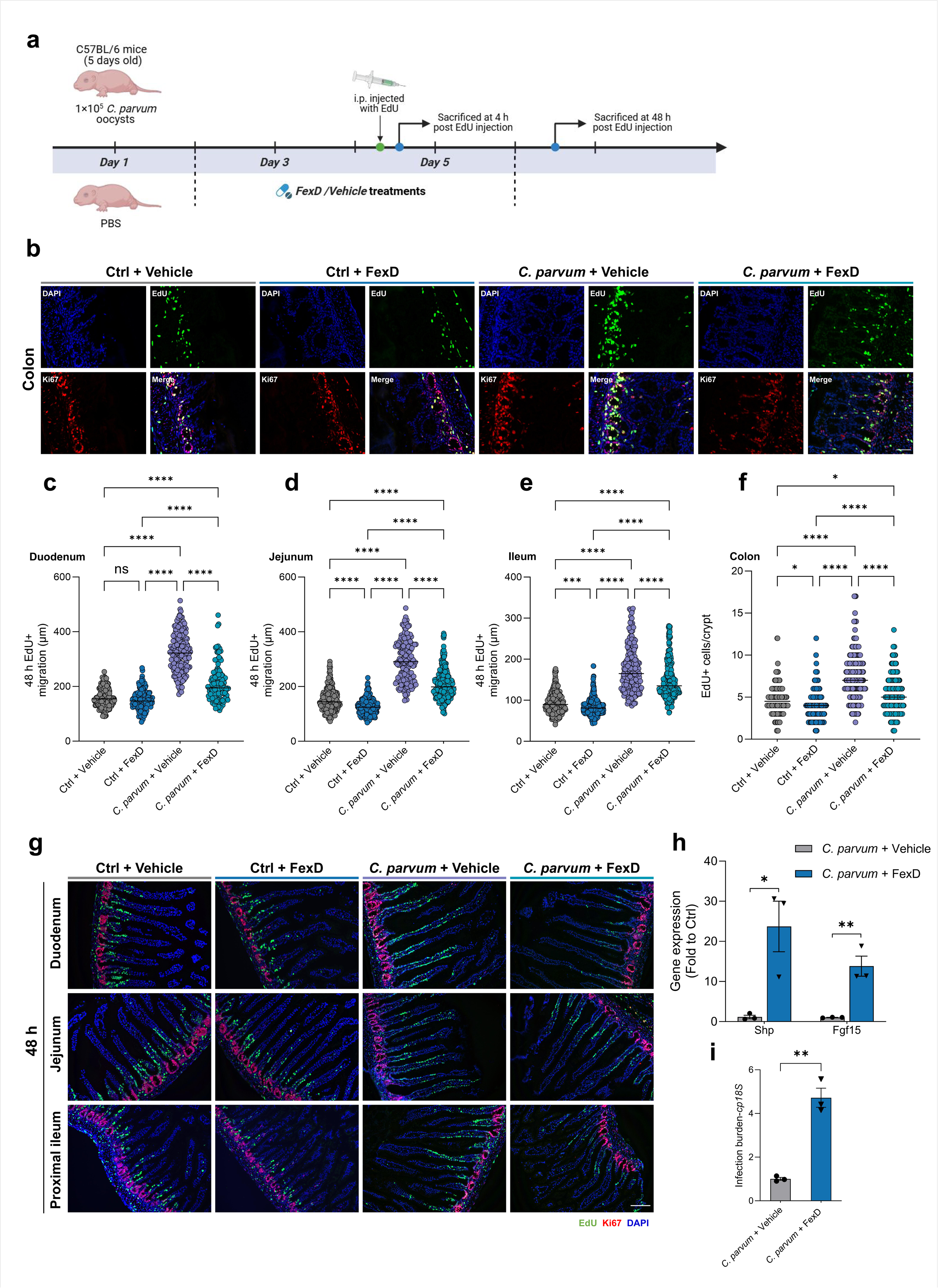
FXR activation attenuates the dysregulated homeostasis in the intestinal epithelium following *C. parvum* infection. **a,** Experiment schematics: Neonatal mice (5 days old) were orally administered 1×10^5^ *C. parvum* oocysts; FexD or vehicle were given at 24 h p.i., followed by daily for 3 days; Mice were i.p. injected with EdU and sacrificed at 4 h post injection; Mice were i.p. injected with EdU, treated with FexD or vehicle for another 2 days and sacrificed at 48 h post injection. **b,** Representative images of Ki67, EdU staining of proliferating cells in crypts at 4 h post injection. Blue: DAPI (nuclei), red: Ki67, green: EdU. Bars, 50 µm. **c-e,** Quantification of cell migration length along the crypt-villus axis at 48 h post injection. Quantification of cell migration length by measuring the distance from the crypt base to the highest labelled cell in the villus (>20 villus per mouse, 3 mice/group). **f,** Quantification of EdU staining of proliferating cells in crypts at 4 h (b) post injection (>50 crypts per mouse, 3 mice/group). **g,** Representative images of Ki67, EdU staining of proliferating cells at 48 h post injection. Blue: DAPI (nuclei), red: Ki67, green: EdU. Bars, 100 µm. **h, i,** Neonatal mice (5 days old) were orally administered 1×10^5^ *C. parvum* oocysts; FexD or vehicle was given at 24 h p.i., followed by daily for 3 days; Ileum tissues were collected, and the mRNA levels of FXR target genes (h) and infection burden (i) were measured by RT-qPCR. Statistical analysis: unpaired t test for (h, i); One-way analysis of variance (ANOVA) with Tukey tests was used for correction of multiple comparisons in c-f, and representative of at least two independent experiments, n = 3 per group. *P < 0.05, **P < 0.01, ***P < 0.001, ****P < 0.0001.

## Discussion

The preferred site of *Cryptosporidium* infection is the ileum, where sporozoites penetrate individual epithelial cells (mainly enterocytes). After cell entry, they form a unique apical localization (intracellular but extra-cytoplasmic), preventing the infection from further spreading to deeper epithelial tissues or entering the circulation and infecting other organs^39,40^. The significant alterations in gene expression profiles and metabolic processes observed in the liver, and proximal small intestine - regions where no parasites were detected - suggest that the pathological changes extend beyond the directly infected ileum. From a pathological perspective, the metabolic dysregulation observed beyond the ileum may play an important role in the growth stunting and malnutrition seen in young children following cryptosporidial infection.

The correlation between lipid and bile acid metabolic dysregulation observed in the liver and intestine from infected mice provides further evidence of the role of the gut-liver axis in the pathogenesis of various enteritis. Given the preferred infection site of *Cryptosporidium* in the ileum, where over 95% of the bile acids are reabsorbed^13^, it maybe pathologically significant that the expression of ASBT is downregulated at the protein level in the infected ileum. This downregulation could reduce bile acid reabsorption, contributing to the decreased bile acid levels in the enterohepatic circulation. Depletion of intestinal cholesterol may also contribute to the decreased bile acid pool, as ASBT function depends on the cholesterol content of lipid raft domains in the plasma membrane^41^. Moreover, the decreased absorptive surface area in the intestine, due to the loss of microvilli and shorten of villus height and intestinal length following *Cryptosporidium* infection, could decrease the brush border apical surface. Surprisingly, the reduction of the bile acid pool and the downregulation of the FXR pathway in the enterohepatic circulation following *C. parvum* infection did not lead to an increase in hepatic bile synthesis as compensation. Instead, there was an overall decrease in bile acid metabolism in the liver. Hepatic bile synthesis generally accounts for only a small fraction of the bile acid pool, as over 95% bile acids are conserved through intestinal reabsorption in the enterohepatic circulation^42^.

While most bile acids are transported back into the liver via enterohepatic circulation, a small fraction of this pool (roughly 5%) escapes reabsorption in the ileum and is further converted into secondary bile acids in the colon by the gut microbiota^13^. Previous studies have demonstrated controversial effects of several bile acids on *Cryptosporidium* infection. Three secondary bile acids (deoxycholate, deoxycholic acid, and lithocholic acid) have been reported to impair *Cryptosporidium* growth *in vitro*^43^. The bile salt sodium taurocholate enhances microneme secretion and increases the entry of *Cryptosporidium* into host cells^34^. Our studies indicate that total bile collected from the gallbladder of adult mice can enhance the entry of *Cryptosporidium* into cultured host epithelial cells. However, adequate supplementation with collected bile significantly reduces the infection burden both *in vivo* and *in vitro*. Additionally, treatment with bile collected from the gallbladder of adult mice or the FXR agonist FexD ameliorates dysregulated cell proliferation and migration in the intestine, including both the directly infected ileum region and the non-infected regions. Several previous studies have suggested a potential association between cryptosporidiosis and the presence of digestive neoplasia^44^. It remains to be determined whether there is an association between Bile acid-FXR pathway and the development of digestive neoplasia following *Cryptosporidium* infection.

Although obvious disturbances in cholesterol and bile homeostasis in the gut-liver axis were observed following *Cryptosporidium* infection, the underlying mechanisms driving these metabolic disturbances remain unclear. In addition to reduced bile acid pool and subsequent lipid malabsorption, the general inflammatory responses triggered by infection as previously reported^45^ may play a role. If so, it would be interesting to explore whether these findings are applicable to infection caused by other gastrointestinal pathogens. In the intestinal epithelium, these disturbances may also reflect the immature state of the epithelial cell population during infection. Previous studies have shown that infection with this parasite accelerates ileal epithelial development, resulting in an epithelium dominated by immature cells^46^. The outgrowth of immature epithelial cells, which are actively progressing through the cell cycle and not yet fully differentiated, may contribute to lipid malabsorption. Biliary infection by *Cryptosporidium*, as reported in AIDS patients and other weakened immune systems^11^, may have additional effects on cholesterol and bile homeostasis. Additionally, bile acids have been shown to drive the maturation of the newborn’s gut microbiota^47^. Gut microbiome development is closely linked to the overall development of the child, with changes in its composition and maturity potentially affecting immune functions, intestinal development, neurodevelopment, nutritional energy harvest, growth, and weight gain^48^. The composition and diversity of the gut microbiome during early development play a crucial role in the incidence and severity of *Cryptosporidium* infection^49–51^. Whether bile treatment influences the maturation of the gut microbiota and, in turn, improves long-term deficits in growth, weight gain, and cognitive development following *Cryptosporidium* infection warrants further exploration.

## Methods

### Ethics statement

All research studies involving the use of animals were reviewed and approved by the Institutional Animal Care and Use Committee (IACUC) of Rush University at Chicago, Illinois, and were carried out in strict accordance with the recommendations in the Guide for the Care and Use of Laboratory Animals.

### *C. parvum* and Cell lines

*C. parvum* oocysts of the Iowa strain were obtained from a commercial source (Bunch Grass Farm, Deary, ID). HCT-8 cells were purchased from ATCC (Manassas, Virginia). The culture media were supplied with 10% fetal bovine serum (FBS) (Ambion, Austin, Texas) and antibiotics (100 IU/ml of penicillin and 100 µg/ml of streptomycin).

### Histology of intestinal tissue

Neonatal mice were killed and small intestine was dissected. Given the stand ’swiss-roll’ was not feasible with the unfixed small intestines of neonatal mice due to their fragility. We collected fresh tissue, gently rolled it into a spiral shape, and embedded it in OCT compound for subsequent analysis. Slides were evaluated in a blinded fashion for measurements of parasite burden, and Oil red staining (see Fig. 5f and Extended Data Fig. 7c). A section four centimeters from the stomach of neonatal mice as the proximal small intestine (jejunum), and the distal small intestine four centimeters from the cecum as the ileum for subsequent analysis (RT-qPCR, Immunoblotting, Immunohistochemistry, and Oil red staining).

### Bile collection

Gall bladders were removed from healthy adult C57BL/6 mice after 24 hours of fasting, punctured with forceps to collect bile, and stored at -80°C until processing.

### *In vitro C. parvum* infection experiments

Models of intestinal cryptosporidiosis using cultured cell lines were employed as previously described^52^. For *C. parvum* infection burden experiments, infection was done in serum-free culture medium for 4 h with a 1:4 ratio of *C. parvum* oocysts to HCT-8 cells. The culture medium was changed to remove free parasites. The medium was supplemented with mouse bile at concentrations of 0.2%, 0.25%, 0.33%, 0.5% and 1% (collected by gall bladder puncture from healthy adult C57BL/6 mice) and cultured for an additional 20 h (a total 24 h p.i.), followed by infection burden measurement. Infection burden was quantified by measuring levels of *C. parvum* 18S gene (*cp18S*)^53^. Immunofluorescent staining of *C. parvum* was carried out using rabbit antiserum against *C. parvum* membrane proteins as previously described^54,55^. For *C. parvum* invasion experiments, HCT-8 cells were incubated with *C. parvum* oocysts (1:4 ratio of oocysts to cells) in the presence of various concentrations of bile (0.07%, 0.2%, 1%, and 2%) or PBS for 4 h, followed by RT-qPCR of *cp18S*.

### *In vivo C. parvum* infection experiments

A well-established infection model of cryptosporidiosis in neonatal mice was used for *in vivo* experiments^56,57^. Neonatal mice (5 days old) were orally inoculated with *C. parvum* (10^6^ oocysts per animal). Mice receiving PBS by oral gavage were used as control. At 72 and 96 h after *C. parvum* oocysts or PBS administration, the animals were sacrificed, and intestine and liver tissues were collected for biochemical analyses. For *C. parvum* infection burden experiments, neonatal mice (5 days old) were orally inoculated with *C. parvum* (at a lower dose of 10^5^ oocysts per animal). At 24 h after *C. parvum* infection, the animals were fed with PBS-diluted bile (0.5, 1.5, 3, and 3,5 µl bile per animal, diluted in PBS to a total volume of 7 μl) and PBS (7µl per animal), followed by daily for 4 days. The animals were sacrificed, and intestine tissues were collected for biochemical analyses.

### RT-qPCR

Total RNA was isolated using TRIzol™ Reagent (Thermo Fisher Scientific, catalogue no. 15596018), and genomic DNA was removed using the DNA-free™ DNA Removal Kit (Thermo Fisher Scientific, catalogue no. AM1906). The reverse-transcription reaction was performed on 2 μg total RNA with M-MLV Reverse Transcriptase (Thermo Fisher Scientific, catalogue no. 28025013). Quantitative PCR assays were conducted using Bio-Rad CFX Manager v3.1 using the iTaq™ Universal SYBR® Green Supermix (Bio-rad, Hercules, CA, USA, catalogue no. 1725124). RNA expression levels were calculated using the ΔΔCt method.

### RNA-Seq and bioinformatic analysis

Total RNAs isolated with Rneasy Mini Kit (QIAGEN, catalogue no. 74104). Three biological replicates (all mice from the same litter) were performed for each condition. The samples were analyzed by BGI Americas Corporation (Cambridge, MA) using the DNBSEQ™ sequencing technology platform, as previously reported^21^. Differential expression analysis and statistical significance were assessed using DESeq in R with a standard negative binomial fit for the aligned counts data and are described relative to the indicated control (Table S1 and S2). Gene set enrichment analysis (GSEA) was performed using the R package “clusterProfiler”^58^ with gene sets from the MsigDB database (http://www.gsea-msigdb.org/gsea/index.jsp)^59^. Gene set variation analysis (GSVA) was conducted with a significance threshold of p < 0.05 to explore correlated pathways, using gene sets from the MsigDB database^60^. Bioinformatic analysis was carried out and results were employed as we previously reported^21,61^.

### Immunoblotting

Membrane and cytosolic protein extracts were isolated from frozen mouse liver or small intestine (30 mg) using the Mem-PER Plus Membrane Protein Extraction Kit (Thermo Fisher Scientific, Catalog no. 89842). Membrane and cytosolic fractions were separated by SDS-PAGE and transferred to a nitrocellulose membrane. Images were captured using the ChemiDoc MP Imaging System (Bio-Rad, Catalog no. 12003154). The following primary and secondary antibodies were used: anti-β Tubulin (Santa Cruz, catalogue no. sc-5274, 1:1000), anti-GAPDH (Santa Cruz, catalogue no. sc-32233, 1:1000), anti-ABCG5 (Thermo Fisher Scientific, catalogue no. BS-5013R, 1:1000), anti-ABCG8 (Novus Biologicals, catalogue no. 1B10A5, 1:1000), anti-SLC10A2 (Abcam, catalogue no. ab203205, 1:1000), anti-FGF15 (Santa Cruz, catalogue no. sc-514647, 1:1000), anti-NR1H4 (Proteintech, catalogue no. 25055-1-AP, 1:3000), anti-NR1H3 (Proteintech, catalogue no. 14351-1-AP, 1:5000), anti-NR1H2 (Proteintech, catalogue no. 60345-1-Ig, 1:1000), anti-Na+/K+-ATPase (Abclonal, catalogue no. A12405, 1:2000), anti-HMGCR (Abclonal, catalogue no. A1633, 1:2000), anti-CYP7A1 (Abclonal, catalogue no. A10615, 1:1000), anti-NR0B2 (Abclonal, catalogue no. A1836, 1:500), anti-HMGCS2 (Abclonal, catalogue no. A19232, 1:2000), anti-rabbit B700 (Bio-Rad, catalogue no. 12004161, 1:2500), anti-mouse B700 (Bio-Rad, catalogue no. 12004158, 1:2500), anti-rabbit B520 (Bio-Rad, catalogue no. 12005869, 1:2500), anti-mouse B520 (Bio-Rad, catalogue no. 12005866, 1:2500), anti-GAPDH hFAB™ Rhodamine (Bio-Rad, catalogue no. 12004167, 1:2500), anti-Tubulin hFAB™ Rhodamine (Bio-Rad, catalogue no. 12004165, 1:2500).

### Immunohistochemistry

For tissue staining, frozen sections (8 μm) were fixed in 4% PFA in PBS for 15 min, then washed twice with PBS. The sections were permeabilized by adding 0.5% Triton X-100 (Bio-rad, catalogue no. 1610407) in PBS for 20 min, followed by removal of the permeabilization buffer and two PBS washes. Sections were then blocked for 30 min in 10% normal donkey serum, 0.1% Triton X-100, 0.01% sodium azide in PBS before incubation with the primary antibody for 2 h at room temperature or overnight at 4 °C. Slides were washed three times with PBS and incubated in block solution containing secondary antibody for 30 min at room temperature. After antibody incubation, sections were washed three times with PBS (5 min per wash) and mounted with Antifade Mounting Medium with DAPI (Vector, catalogue no. H-1200). Images were captured using a Keyence BZ-X810 microscope. For cell staining, HCT-8 cells were fixed in 4% PFA in PBS for 15 min and washed twice with PBS. Fixed cells were permeabilized with 0.1% Triton X-100 in PBS for 5 min and washed three times with PBS. Cells were blocked for 30 min in 10% normal donkey serum, 0.1% Triton X-100, 0.01% sodium azide in PBS, followed by incubation with primary antibody for 1 h at room temperature. Cells were then washed three times with PBS and incubated in block solution containing secondary antibody for 30 min at room temperature.

Following antibody incubation, cells were washed three times with PBS (5 min per wash) and mounted with Antifade Mounting Medium with DAPI (Vector, catalogue no. H-1200). Images were taken using a Keyence BZ-X810 microscope. Immunofluorescent staining of *C. parvum* was carried out using rabbit antiserum against *C. parvum* membrane proteins as previously described^54,55^. Other antibodies used for immunofluorescent staining include anti-Ki67 antibody (Cell Signaling Technology, catalogue no. 9129, 1:200), anti-ABCA1 antibody (Thermo Fisher Scientific, catalogue no. PA1-16789, 1:200), goat-anti-bovine Alexa fluor 488 (Jackson Immuno Research, catalogue no. 101-545-003, 1:250), goat-anti-bovine Alexa fluor 594 (Jackson Immuno Research, catalogue no. 101-585-003, 1:250), donkey-anti-rabbit Alexa fluor 488 (Jackson Immuno Research, catalogue no. 711-545-152, 1:500), donkey-anti-rabbit Rhodamine Red (Jackson Immuno Research, catalogue no. 711-295-152, 1:500).

### Oil red staining

Lipid vesicle storage in tissue was assessed with Oil Red O staining, following the manufacturer’s protocol (Abcam, catalogue no. ab150678). Images were captured using a Keyence BZ-X810 microscope.

### Cell viability

Cell viability was assessed using a Neutral Red Assay Kit, following the manufacturer’s instructions (Abcam, catalogue no. ab234039). The absorbance of each well at 540 nm was measured using a microplate reader (SpectraMax iD3, Molecular Devices, USA) to evaluate the cell viability.

### Bile Acid Targeted Metabolomics Assay

Metabolites were extracted from the livers and small intestines of control and *C. parvum*-infected neonatal mice, with three biological replicates for each condition. The samples were analyzed by Metware Biotechnology Inc (Metware). Metabolomics profiling was employed as previously described^62^.

### Quantitative Lipidomics

Metabolites were extracted from the livers and proximal intestinal epithelium of control and *C. parvum* infected neonatal mice, with four biological replicates for each condition. The samples were analyzed by Metware Biotechnology Inc (Metware). Metabolomics profiling was employed as previously described^63^.

### Cholesterol assay

Cholesterol Quantification Assay Kit (Sigma-Aldrich, CS0005) was used to measure the cholesterol concentration of samples. We sampled the distal small intestine four centimeters from the cecum of neonatal mice and collected epithelial cells. Briefly, 46 μL Assay Buffer, 2 μL Probe, 2 μL Enzyme Mix, and 50 μL sample were mixed and incubated at 37 °C for 30 min in each well. A calibration curve was established for every measurement with standard samples with 0–5 μg cholesterol. All samples were diluted to the range of the calibration curve with the Assay Buffer. Absorbance at 570 nm was measured and compared to the standards on the same plate to determine total cholesterol.

### *In vivo* EdU assay

Neonatal mice (5 days old) were orally inoculated with *C. parvum* (10^5^ oocysts per animal). Mice receiving PBS by oral gavage were used as control. For bile treatment, 24 h after *C. parvum* oocysts or PBS administration, control mice were treated with PBS, *C. parvum*-infected mice were fed PBS-diluted bile (7µl per animal,1:1 ratio of bile to PBS) or PBS only (7µl per animal), followed by daily treatments for 3 days. Mice were i.p. injected with EdU and sacrificed 4 h post injection. In another group, mice were i.p. injected with EdU, treated with PBS-diluted bile or PBS for an additional 2 days, and sacrificed 48 h post injection. For FXR agonists fexaramine D (FexD) treatment, 24 h after *C. parvum* oocysts or PBS administration, control mice were i.p. injected with vehicle, and *C. parvum*-infected mice were i.p. injected with vehicle with or without 100 mg/kg of FexD, followed by daily treatments for 3 days. Mice were i.p. injected with EdU and sacrificed 4 h post injection, or were i.p. injected with EdU, treated with vehicle with or without FexD for an additional 2 days and sacrificed 48 h post injection. Intestine tissues were frozen in OCT and cryosectioned (10 µm). EdU staining was performed using the Click-iT EdU imaging kit (Thermofisher cat# C10337) according to the manufacturer’s protocol. To evaluate the proliferation and migration of EdU-labeled cells, we calculated the average values across all villi/crypts within the same group as previously reported^64^.

### Bioinformatic analysis of metabolomics

Multivariate analysis of Variable Importance in Projection (VIP) from OPLS-DA modeling (MetaboAnalystR, v.1.0.1) was used to preliminarily select differential metabolites from different samples. For two-group analysis, differential metabolites were determined by VIP (VIP > 1) and P-value (P-value < 0.05, Student’s t test). The data were log-transformed (log_2_) and mean-centered prior to OPLS-DA. To avoid overfitting, a permutation test (200 permutations) was performed. For KEGG annotation and enrichment analysis, identified lipids were annotated using the KEGG Compound database (http://www.kegg.jp/kegg/compound/). Annotated lipids were then mapped to the KEGG Pathway database (http://www.kegg.jp/kegg/pathway.html). Pathways with significantly regulated lipids were further analyzed using the MSEA (lipid sets enrichment analysis), with significance determined by the hypergeometric test’s p-values.

### Statistical analysis

For all quantitative analyses, a minimum of three biological replicates were included. Statistical tests were selected based on the assumption that sample data are from a population following a probability distribution with a fixed set of parameters. Student’s t-tests were used to assess the statistical significance of differences between two groups. One-way AVOVA was used for multiple comparison tests. The following values were considered statistically significant: *P < 0.05, **P < 0.01, ***P < 0.001, ****P < 0.0001. Calculations were performed using the GraphPad Prism 10 software package. Data are presented as mean ± the standard error of the mean.

### Data availability

The RNA-seq data generated in this study have been deposited into NCBI’s Gene Expression Omnibus (GEO) and are accessible through GEO series accession number GSE234579 and GSE272799. The metabolomics analysis data are presented in Table S3-S6. Uncropped images of immunoblots presented in the figures are included in Supplementary Figure 1. Source data are available with this paper.

## Supplemental Figures Legends

**Extended Data Fig.1.**
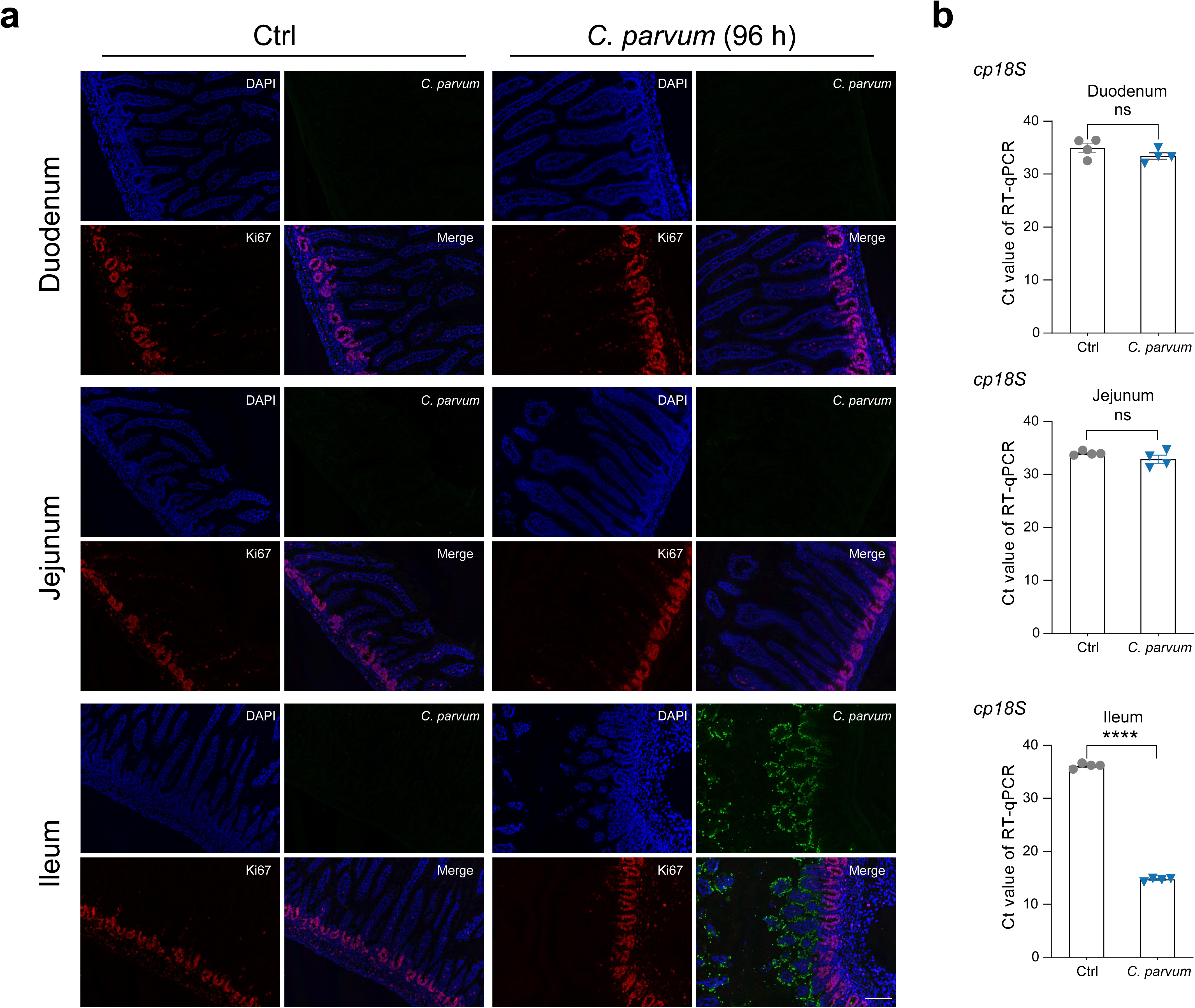
*C. parvum* intestinal infection burden in neonates with and without *C. parvum* infection. **a,** Representative images of duodenum, jejunum and ileum sections from *C. parvum* infected and control mice immunostained for *C. parvum* and Ki67 (96 h p.i.). Blue: DAPI (nuclei), red: Ki67, green: *C. parvum*. Bars, 100 µm. **b,** Relative amounts of mRNAs analysed by RT–qPCR. Neonatal mice (5 days old) were inoculated with *C. parvum* oocysts (10^6^ oocysts per animal) for 96 h. Infection burden was evaluated by RT-qPCR of *cp18S* gene in the duodenum, jejunum and ileum, respectively. The cycle threshold (Ct) values were used to determine the infection burden. Each symbol represents one mouse, all from the same litter. Statistical analysis: unpaired t test for b. Data is expressed as mean ± SEM, and representative of at least two independent experiments, n = 4 per group. *P < 0.05, **P < 0.01, ***P < 0.001, ****P < 0.0001.

**Extended Data Fig.2.**
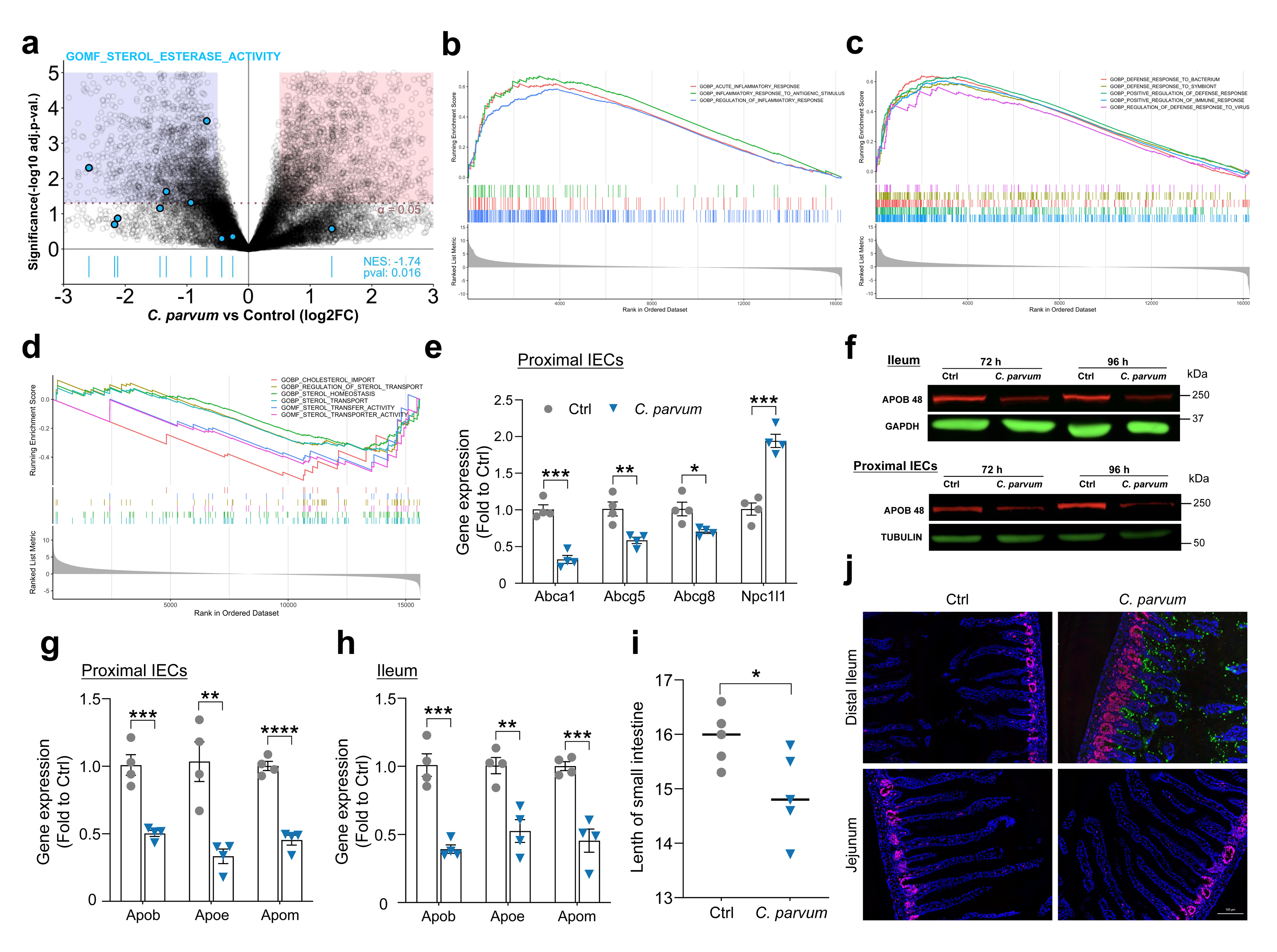
*C. parvum* infection disrupts cholesterol homeostasis in the small intestine. **a,** Differential gene expression analysis upon *C. parvum* infection (72 h p.i.). The enrichment of sterol esterase activity gene set is shown at the bottom. **b-d,** GSEA showing the enriched pathways of inflammatory reactions (b), defense response (c), sterol homeostasis and sterol transport (d) (|NES| > 1, p < 0.05, FDR < 0.25). **e, g, h,** Relative amounts of mRNAs analyzed by RT–qPCR. Gene expression was presented as fold change to uninfected control. **f,** Immunoblotting analysis of proximal and ileal IECs samples. GAPDH and TUBULIN were used as loading control of cytosolic protein. **i,** Small intestine length of control and *C. parvum* infected mice (120 h p.i.). **j,** Sections from the proximal/Jejunum and distal small intestine of neonatal mice 4 days after *C. parvum* infection were immunostaining for Ki67 and *C. parvum*. Blue: DAPI (nuclei), red: Ki67, green: *C. parvum*. Each symbol represents one mouse, all from the same litter. Statistical analysis: unpaired t test for (e, g-i). Data are expressed as mean ± SEM, and representative of at least two independent experiments, n = 3–5 per group. *P < 0.05, **P < 0.01, ***P < 0.001, ****P < 0.0001. For gel source data, see Supplementary Figure 1.

**Extended Data Fig.3.**
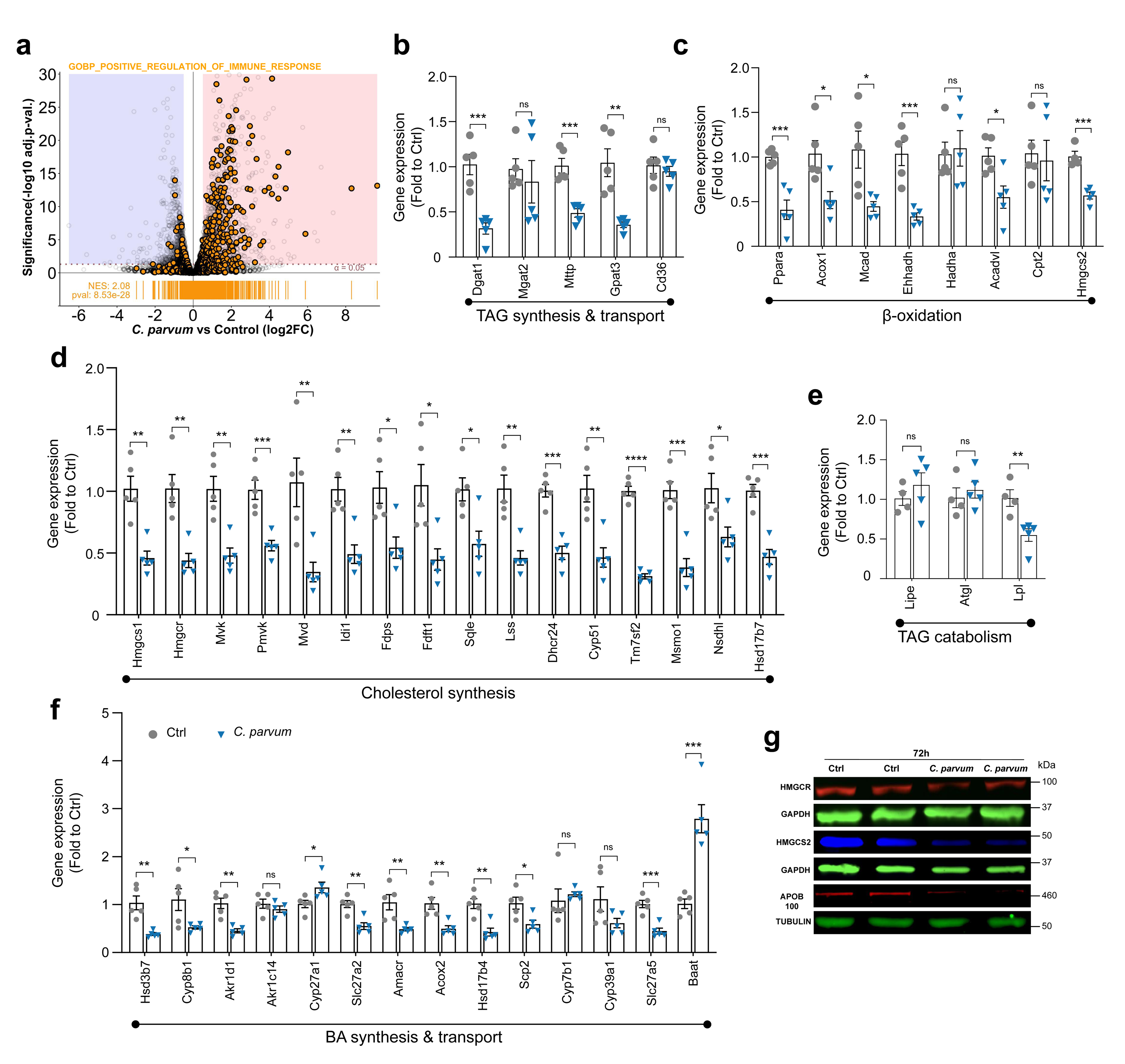
*C. parvum* infection alters hepatic genes expression. **a,** Differential hepatic gene expression analysis upon *C. parvum* infection (72 h p.i.). The enrichment of positive regulation of immune response gene set is shown at the bottom. **b-f,** Relative mRNA expression of lipid metabolic genes associated with the indicated processes were measured by RT-qPCR. **g,** Immunoblotting analysis of liver samples in neonates with and without *C. parvum* infection (72 h p.i.). GAPDH and TUBULIN were used as loading control of cytosolic protein. Each symbol represents one mouse, all from the same litter. Statistical analysis: unpaired t test for (b-f). Data are expressed as mean ± SEM, and representative of at least two independent experiments, n = 4–5 per group. *P < 0.05, **P < 0.01, ***P < 0.001, ****P < 0.0001. For gel source data, see Supplementary Figure 1.

**Extended Data Fig.4.**
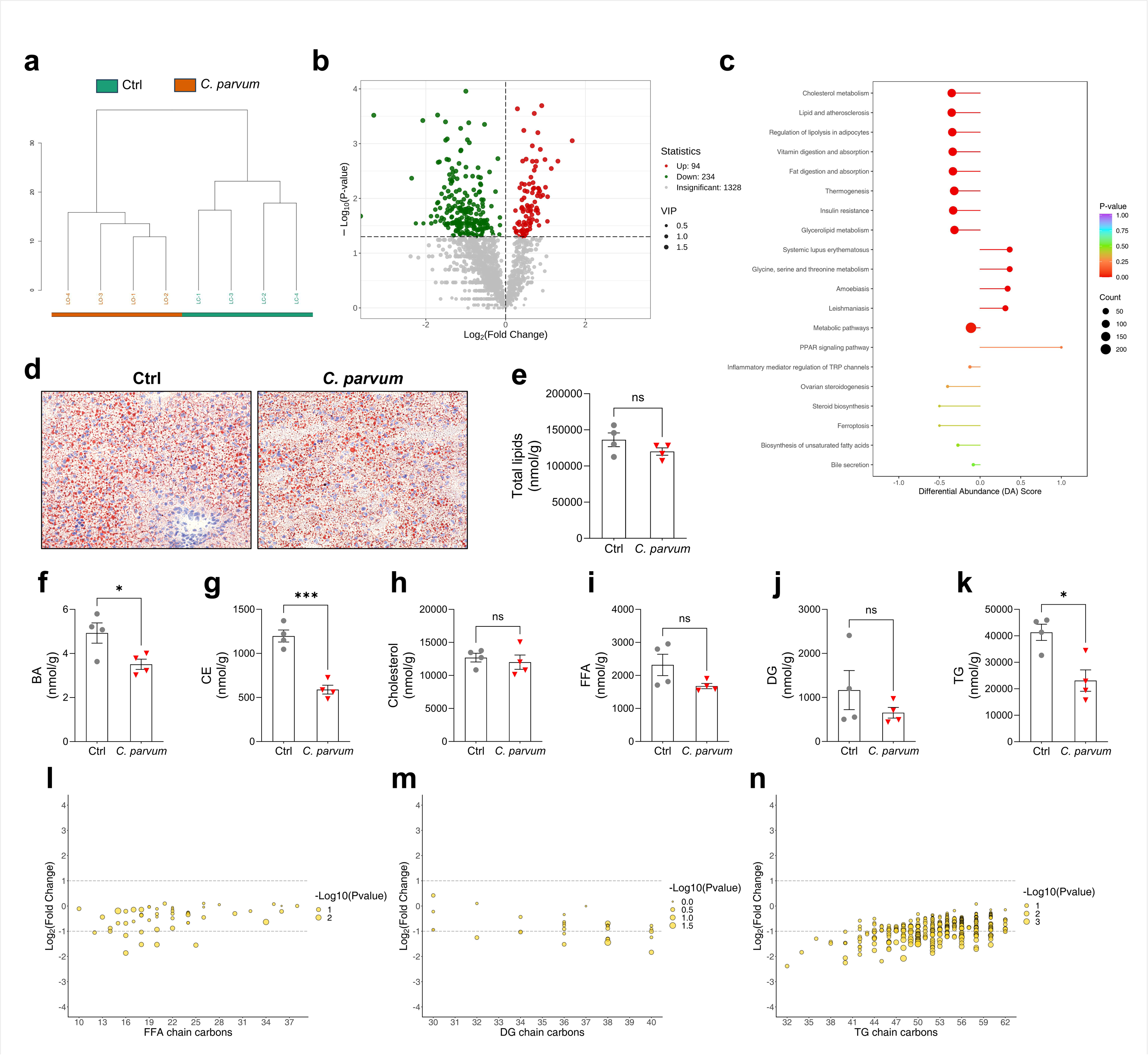
Impaired lipid homeostasis in the livers of *C. parvum* infected mice. **a,** Hierarchical clustering showing the similarity between control and *C. parvum* infected neonatal mice (96 h p.i.). **b,** Volcano plot illustrating the hepatic lipids expression between control and *C. parvum* infected neonatal mice (96 h p.i.). Lipid with VIP ≥ 1 represents significant difference. **c,** KEGG pathway enrichment analysis and differential abundance (DA) scores of differential metabolites in the liver of control and *C. parvum* infected neonatal mice (96 h p.i.). DA score = (up regulated lipids in a pathway-down regulated lipids in a pathway) / (Total number of lipids annotation in a pathway). **d,** Oil Red O stains of the liver sections in neonates with and without *C. parvum* infection (96 h p.i.). **e,** The total quantification of all hepatic lipids between control and *C. parvum* infected neonatal mice (96 h p.i.). **f-k,** Graphs depicting quantification of the differences in hepatic BA (f), CE (g), Cholesterol (h), FFA (i), DG (j) and TG (k) between control or *C. parvum* infected mice (96 h p.i.). **l-n,** Graphs showing the fold change of hepatic FFA (l), DG (m) and TG (n) with different chain lengths between control or *C. parvum* infected mice (96 h p.i.). Each point represents a lipid, the size of the point represents the P-value. Each symbol represents one mouse, all from the same litter. Statistical analysis: unpaired t test for (e-k). Data are expressed as mean ± SEM, n = 4 per group. *P < 0.05, **P < 0.01, ***P < 0.001, ****P < 0.0001.

**Extended Data Fig.5.**
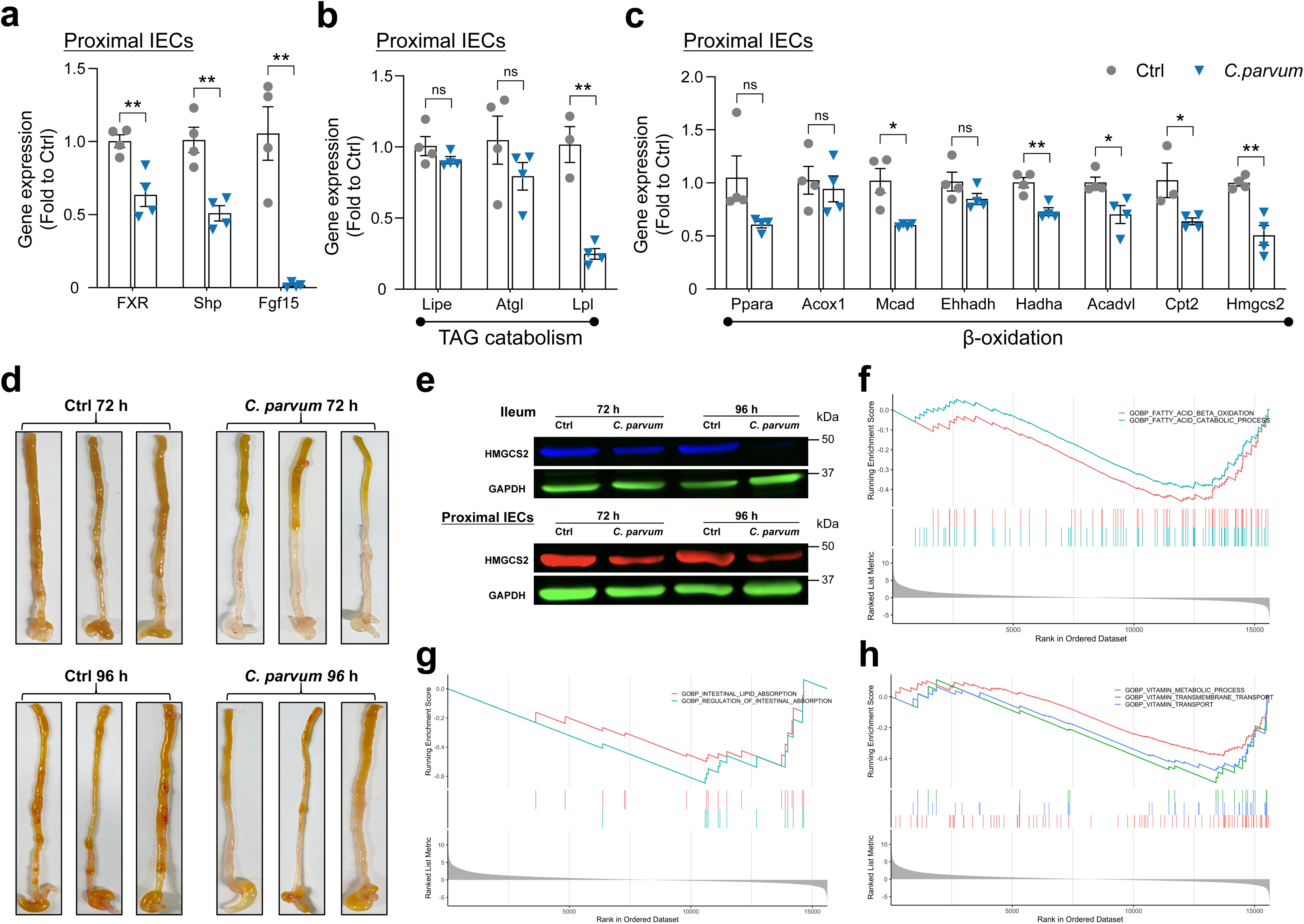
*C. parvum* infection alters intestinal lipid metabolic genes expression. **a-c,** Relative amounts of mRNAs analyzed by RT–qPCR. Gene expression was presented as fold change to uninfected control. **d,** Representative images of ileum (72 h and 96 h p.i.). **e,** Immunoblot analysis of proximal and ileal IECs from control and *C. parvum* infected neonatal mice (72 h and 96 h p.i.). GAPDH was used as loading control of cytosolic protein. **f-h,** GSEA showing the enriched pathways of fatty acid metabolic process (f), intestinal lipid absorption process (g) and vitamin metabolic process (h) (|NES| > 1, p < 0.05, FDR < 0.25). Each symbol represents one mouse, all from the same litter. Statistical analysis: unpaired t test for (a-c). Data are expressed as mean ± SEM, and representative of at least two independent experiments, n = 3–4 per group. *P < 0.05, **P < 0.01, ***P < 0.001, ****P < 0.0001. For gel source data, see Supplementary Figure 1.

**Extended Data Fig.6.**
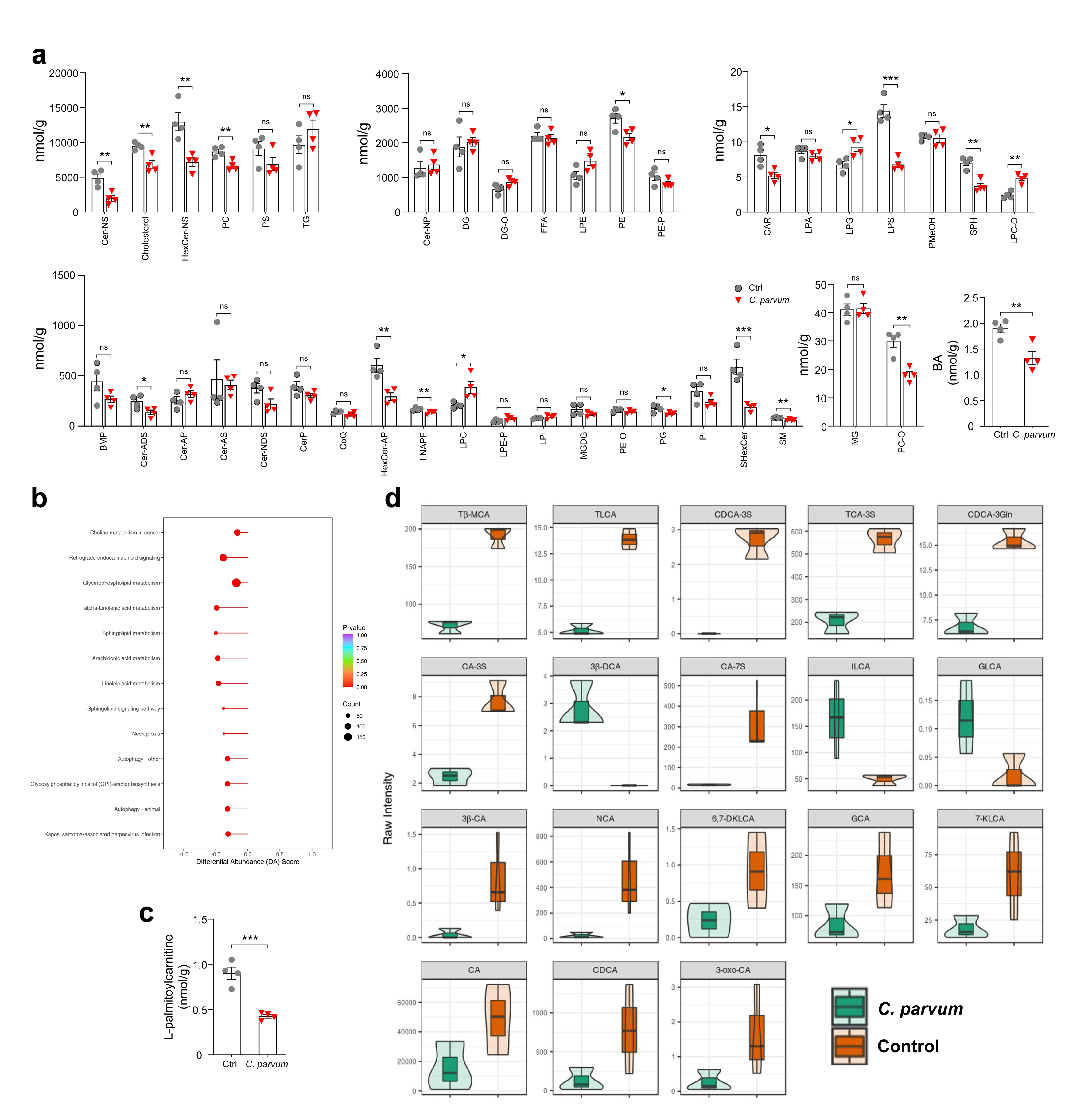
Impaired lipid homeostasis in the proximal IECs of *C. parvum* infected mice. **a,** Graphs depicting quantification of the differences in proximal IECs lipid abundance of each subclass between control or *C. parvum* infected mice (96 h p.i.). **b,** KEGG pathway enrichment analysis and DA scores of differential metabolites in proximal IECs of control and *C. parvum* infected neonatal mice (96 h p.i.). **c,** Graphs showing the quantification of L-palmitoylcarnitine in proximal IECs between control or *C. parvum* infected mice (96 h p.i.). **d,** Violin plot showing the result of top 18 differentially bile acids with the largest log2FC value. The box in the middle represents the interquartile range, and the middle box represents the 95% confidence interval. The black horizontal line is the median, and the outer shape represents the distribution density of the data. Each symbol represents one mouse, all from the same litter. Statistical analysis: unpaired t test for (a,c). Data are expressed as mean ± SEM, and representative of at least two independent experiments, n = 3–4 per group. *P < 0.05, **P < 0.01, ***P < 0.001, ****P < 0.0001.

**Extended Data Fig.7.**
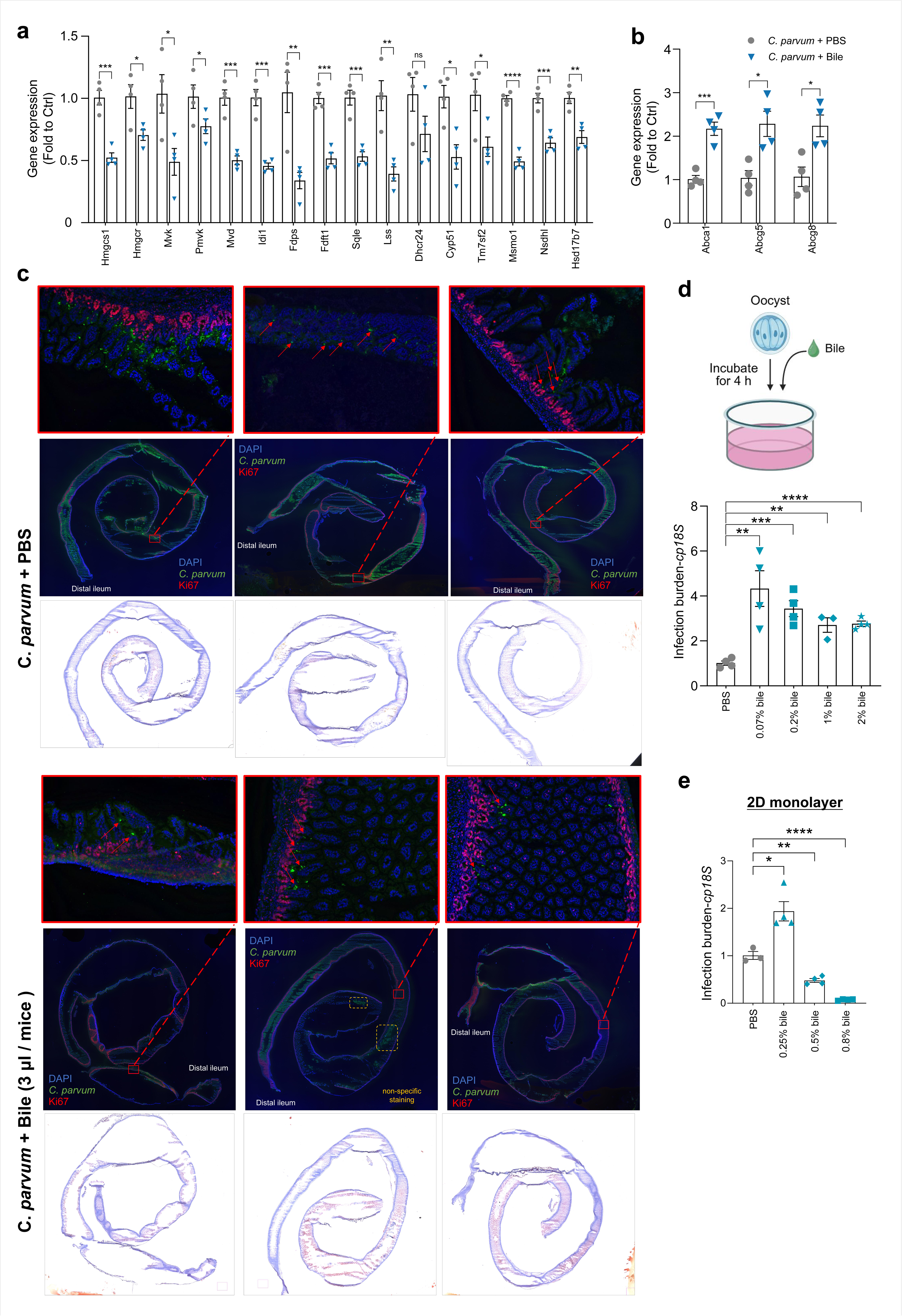
Effects of bile on *C. parvum* infection. **a,** Effects of bile on *C. parvum* intestinal infection in neonates. Neonatal mice (5 days old) were orally administered *C. parvum* oocysts; PBS or bile were given at 24 h p.i. followed by daily for 4 days, respectively; Intestinal tissues were collected, and the mRNA levels of cholesterol biosynthesis genes (a) and cholesterol efflux genes (b) were measured by RT-qPCR. Gene expression was presented as fold change to uninfected control. **c,** Immunofluorescent and Oil Red O staining of the distal small intestine sections from (a). A higher magnification of the boxed region (red) to visualize the *C. parvum* staining; Red arrow indicates *C. parvum* staining. Blue: DAPI, red: Ki67, green: *C. parvum*. **d,** Effects of bile on *C. parvum* invasion of cultured HCT-8 cells. Cells were incubated with oocysts for 4 h in the presence of different concentrations of bile or PBS, followed by RT-qPCR of *cp18S*. **e,** Effects of bile on *C. parvum* infection of 2D monolayers. Cells were incubated with oocysts for 4 h first, subsequently exposed to different concentration of bile or PBS for another 20 h, followed by RT-qPCR of *cp18S*. Each symbol represents one mouse, all from the same litter. Statistical analysis: unpaired t test for (a, b, d, e). Data are expressed as mean ± SEM, and representative of at least two independent experiments, n = 4 per group. *P < 0.05, **P < 0.01, ***P < 0.001, ****P < 0.0001.

**Extended Data Fig.8.**
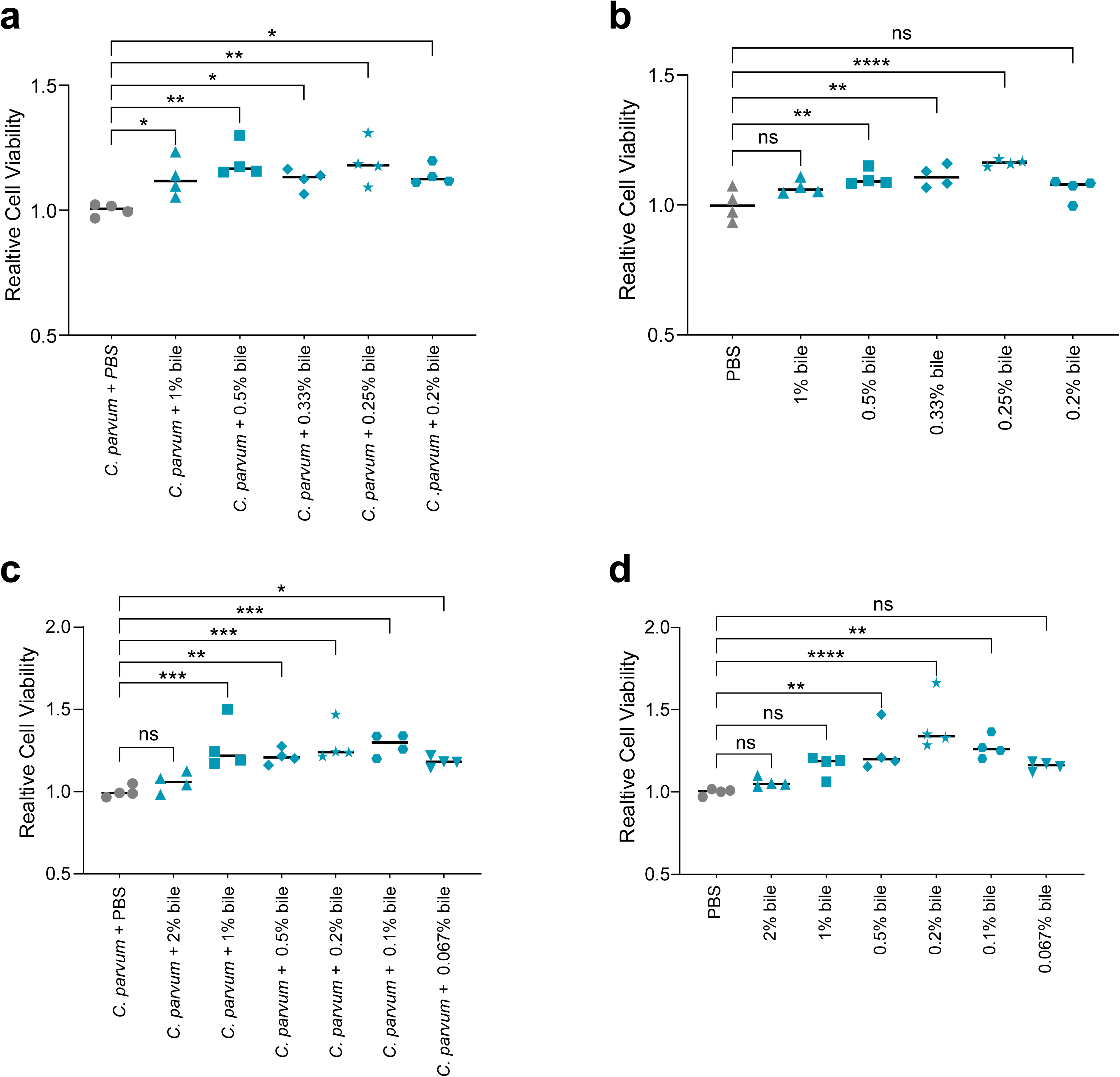
Effects of bile on HCT-8 cells viability with or without *C. parvum* infection. **a,** Effects of bile on HCT-8 cells viability in the presence of *C. parvum* infection. Cells were incubated with oocysts for 4 h first, subsequently exposed to different concentrations of bile or PBS for another 20 h, followed by cell viability assay. **b,** Cell survival of HCT-8 cells treated with different concentration of bile or PBS for 24 h. (c, d) Cell survival of HCT-8 cells exposed to different concentration of bile or PBS with (c) or without (d) oocysts for 4 h. One-way analysis of variance (ANOVA) with Dunnett’s multiple comparison test was used for correction of multiple comparisons in a-d, n = 4 per group. *P < 0.05, **P < 0.01, ***P < 0.001, ****P < 0.0001.

**Extended Data Fig.9.**
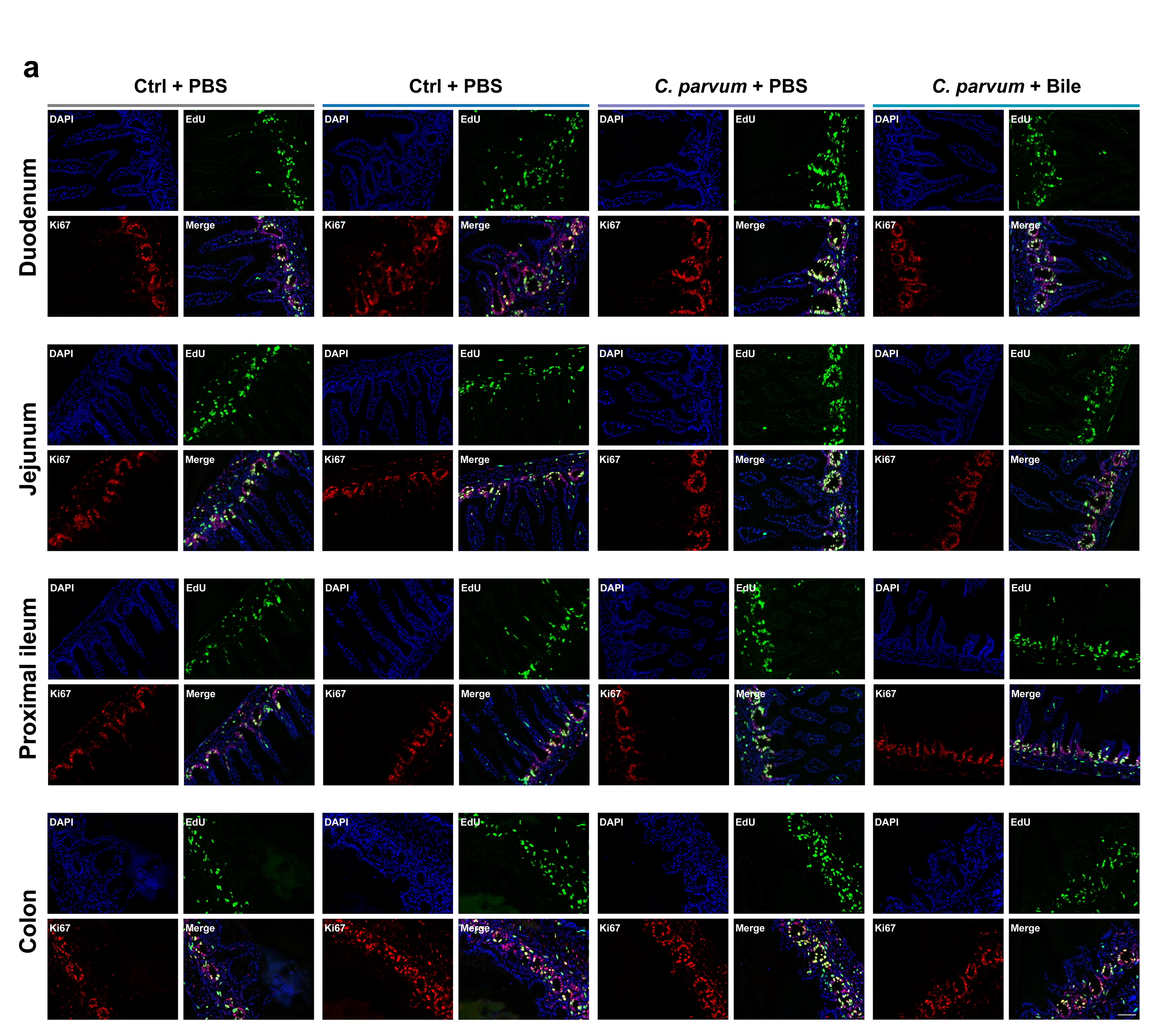
Bile intervention attenuates the dysregulated cell proliferation in crypts following *C. parvum* infection. **a,** Neonatal mice (5 days old) were orally administered 1×10^5^ *C. parvum* oocysts; PBS or bile was given at 24 h p.i. followed by daily for 3 days; Mice were i.p. injected with EdU and sacrificed at 4 h post injection. Representative images of Ki67, EdU staining of proliferating cells in crypts at 4 h post injection. Blue: DAPI (nuclei), red: Ki67, green: EdU. Bars, 50 µm.

